# Adaptation in the eye and brain contributes to species divergence in visual perception in *Heliconius* butterflies

**DOI:** 10.64898/2026.08.27.747543

**Authors:** Daniel Shane Wright, José Borrero, Yi Peng Toh, Lisa Ammer, Anupama Nayak Manel, J. Benito Wainwright, Juanita Gutiérrez-Valencia, Lucie Queste, Elisa Mogollon Perez, Michelle Guachamin-Rosero, Pamela Chamba-Vaca, Daniela Lozano-Urrego, Geraldine Rueda-Muñoz, Patricio A. Salazar-Carrion, Nicola J. Nadeau, Chris D. Jiggins, Carolina Pardo-Diaz, Camilo Salazar, Caroline N. Bacquet, Stephen H. Montgomery, Richard M. Merrill

## Abstract

Sensory systems mediate the interaction between organisms and their environment, but how complex sensory pathways evolve and relate to variation in perception and behavior across ecological contexts, remains poorly understood, especially for terrestrial taxa. Here, we investigate whole-visual-system adaptation in *Heliconius erato* butterflies. Using continent-wide sampling, we demonstrate that within *H. erato*, facet count significantly decreased with increasing elevation. Common-garden rearing of low-elevation *H. erato* populations from Ecuador and their high-elevation sister species, *H. himera*, showed that eye and brain morphology are heritable, and comparisons to genomic measures of divergence indicates that this variation is due to divergent selection. Parallel comparisons from Colombia involving *H. chestertonii* (high elevation) and *H. erato venus* (low elevation) further revealed that eye and brain morphology can evolve as independent, decoupled traits. For both locations, differences in visual acuity correlated with variation in facet count. We also observed parallel evolution of spectral sensitivity, with independent high-elevation populations having fewer red-reflecting lateral filtering pigments. To experimentally link visual system morphology to behavior, we assessed visual acuity in second-generation *H. erato cyrbia*-*H. himera* hybrids. Overall, acuity was influenced by facet count, and when analyzed together with brain morphology, by a positive interaction between facet count and optic lobe volume, demonstrating that structural investment in the eye and neural expansion combine to maximize visual perception. This work shows that visual adaptation is a multi-layered process whereby sensory traits can evolve independently under localized ecological pressures, but evolution across the visual pathway contributes to refinements in behavioral performance.

**Significance statement:** By integrating broad surveys of wild populations, common-garden rearing, and behavioral assays in *Heliconius* butterflies, we show that visual adaptation along elevational gradients is a multi-layered process. Eye morphology, neuroanatomy, and spectral sensitivity display independent, habitat-associated shifts, with peripheral and neural components specifically evolving under divergent selection. Crucially, behavioral tests in hybrid crosses reveal that eye and brain traits interactively influence visual acuity, showing that neural expansion must accompany peripheral changes to improve visual performance. This modular yet co-evolving response across the visual pathway provides a flexible mechanism for sensory adaptation across terrestrial environments.

## Introduction

Understanding how populations adapt and diverge is a central theme of evolutionary biology. While focus often falls on morphological traits, the interface between any individual and its environment is mediated by its sensory systems. In many animals, the visual system plays an integral role in dictating how individuals perceive and interact with their surroundings, directly shaping both survival and reproductive success. Consequently, natural selection is expected to favor divergent visual traits in populations exposed to different sensory conditions, or which exploit different resources. Habitat-associated variation in visual traits is well-documented in aquatic taxa, influencing both local adaptation and species divergence (1, 2). However, although similar processes are expected on land (3), sensory conditions are more variable than in water, and fewer examples of visual and behavioral divergence between terrestrial species are known (3). Furthermore, how selection shapes adaptation across different levels of the visual pathway is rarely investigated (but see (4, 5)), and formal links to changes in behavior limit clarity over the behavioural importance of case studies of sensory evolution. Addressing both these limitations is crucial to fully appreciate how visual adaptations track changes in local ecology, and how this may contribute to species divergence.

Neotropical *Heliconius* butterflies offer an excellent opportunity to determine how terrestrial environments may shape different components of the visual pathway. These butterflies are found throughout tropical and subtropical regions of the Americas, inhabiting variable forest habitats (6), and are well known for their mimetic warning patterns. Because these patterns experience strong divergent selection due to predation (7, 8), and are also used during mate choice (9–13), their divergence has been considered a key step during *Heliconius* speciation (14). However, the maintenance of within-species geographic morphs suggests that color pattern divergence must be coupled with additional adaptations and/or ecological transitions to permit species divergence.

*Heliconius* rely on vision for a number of key ecological tasks, including foraging and host plant selection (15, 16), in addition to mate choice. As in all insects, visual signal integration begins with the compound eye that is comprised of numerous independent photosensitive units (ommatidia) that receive light information and transfer it to the brain. While opsin expression shows primarily clade-level shifts, diverging between lineages separated by ∼5-10my and remaining more conserved at shallower phylogenetic scales (17–19), eye morphology in *Heliconius* appears to differ between closely related species inhabiting different forest types. For example, compared to *H. melpomene*, which is found at the forest edge, the closed canopy specialist, *H. cydno*, has larger eyes with more facets (the hexagonal lens of each ommatidia) (20–22). These differences mirror adaptations within the brain, as *H. cydno* also has increased investment in several visual neuropils compared to *H. melpomene* (20, 23). This suggests that selection is distributed across the visual pathway, but whether the visual system evolves as a unified system or as a mosaic of distinct adaptations, and how these vary more broadly across habitat types both within and between species, is unresolved.

Here, we use an integrative approach to test how whole visual system adaptation may influence behavioral evolution and species divergence within the *H. erato* species complex. *H. erato* populations occur widely from sea level to ∼1600m (24), an elevational gradient characterized by shifts in canopy height, gap density, and understory light availability (25), and previously linked to genomic and phenotypic divergence (26–31). In particular, two high-elevation species within the *erato* group, *H. himera* and *H. chestertonii*, have reduced investment in several visual neuropils compared to parapatric lower-elevation *H. erato* populations (32, 33). Building on this, we generate eye morphology data from wild *H. erato* populations spanning Central and South America to test for general effects of elevation. We then explore whether divergence in multiple visual system traits is produced by adaptive processes, and test how these traits together predict variation in visual acuity. Collectively, our data show that visual adaptation in *Heliconius* is a multi-layered process whereby different traits can evolve independently, but coordination across components is required to increase visual acuity. Such modular sensory architectures provide a highly responsive pathway for generating ecological divergence.

## Results

### Elevation predicts variation in facet count

To begin to investigate the evolution of the visual system in *H. erato* and its related species, we quantified eye morphology of 314 wild butterflies from 32 sampling locations (Fig. 1A, B; Table S1). Facet count was strongly predictive of overall corneal area across all populations (R^2^ = 0.81, p <0.001, Fig. 1C), suggesting that eye size variation within the *erato* species complex is primarily driven by changes in facet count, rather than facet size, with one exception, *H. erato venus* (see below; otherwise analyses of corneal area were quantitatively similar and are presented in supplementary information). For all analyses presented below, we always account for sex and an allometric control (i.e. tibia length or central brain volume) unless otherwise specified (see methods for full model descriptions).

**Figure 1.**
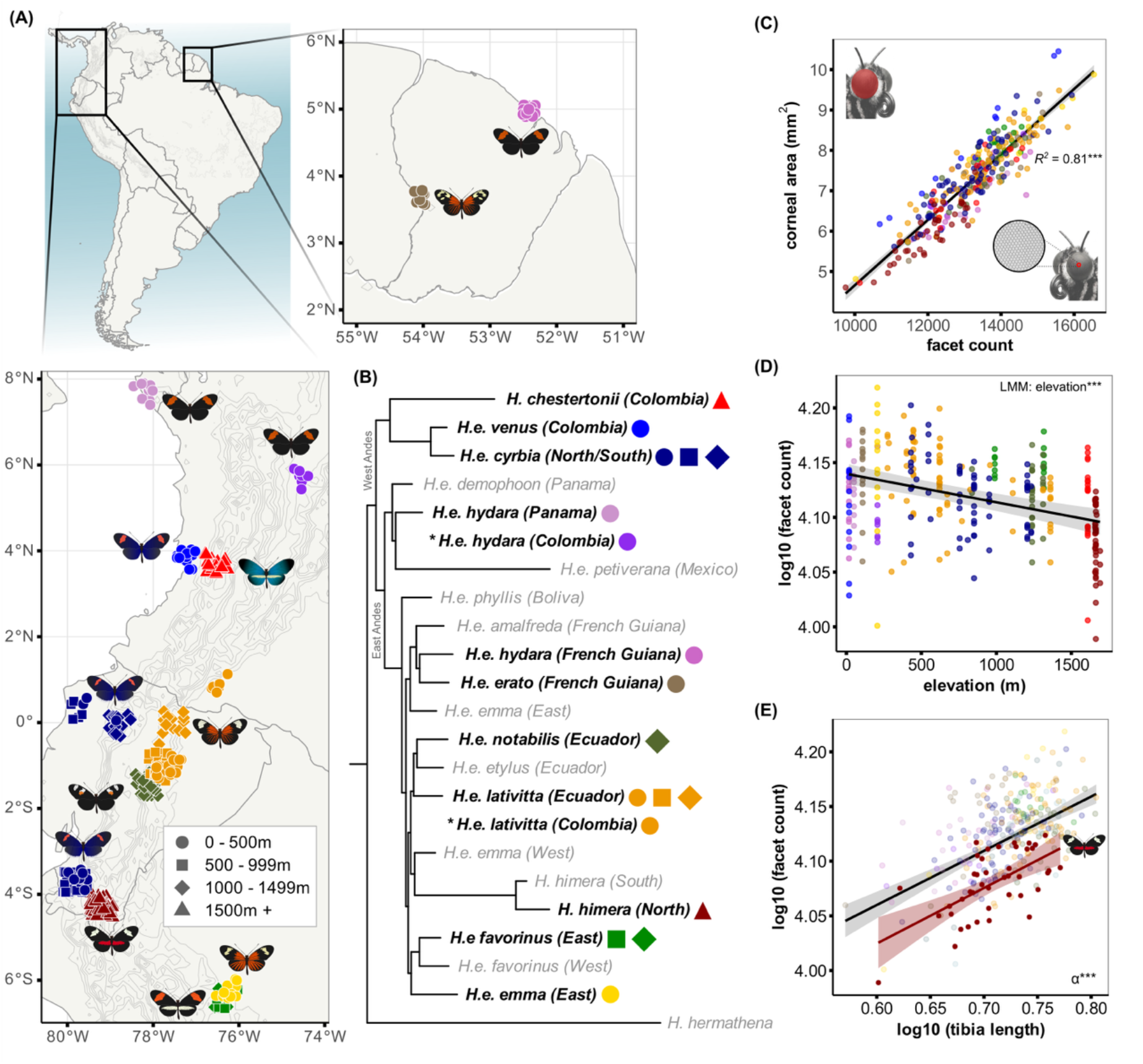
Elevation predicts variation in facet count in *H. erato*. We quantified eye morphology for **(A)** multiple wild *H. erato* populations. **(B)** Phylogenic tree recreated from Van Belleghem et al. (34); \**H. erato hydara* and \**H. erato lativitta* from Colombia were not included in the previous tree, and populations in grey were not included in this study. **(C)** Total facet count and corneal area were highly correlated, suggesting that variation in overall eye size is due to differences in facet count. Shaded ribbon represents standard error. (**D)** Across all samples, higher elevations were associated with fewer facets. The solid black represents the estimated marginal mean and 95% confidence interval (shaded ribbon) from a model accounting for sex, tibia length, and population (as a random effect). **(E)** For the high-elevation specialist *H. himera*, there was a significant grade shift (α) in the relationship between facet count and tibia length versus the other populations, indicating reduced facet count for *H. himera*. The shaded ribbons represent standard error. LMM = linear mixed model; ***indicates p <0.001.

We observed between-population variation in total facet count (linear mixed model [LMM]: Wald χ^2^ = 52.79, p <0.001; Fig. S1) but little evidence of phylogenetic structure, including between eastern and western clades which represent the major phylogenetic split within *H. erato* (0.8–1.0 mya; (34); Fig. 1B). Previous comparisons reported reduced investment in several visual neuropils for the high-elevation specialists *H. himera* and *H. chestertonii* compared to their lower elevation counterparts (32, 33), and here, our broad-scale sampling revealed a continuous negative effect of elevation on facet count (Fig. 1D). The elevation effect remained regardless of whether the analysis included the high-elevation nested specialists, *H. himera* and *H. chestertonii* (LMM: Wald χ^2^ = 38.69, p <0.001; Fig. 1D), or was restricted to only *H. erato sensu stricto* (LMM: Wald χ^2^ = 23.43, p <0.001; Fig. S2). Compared to the other sampled populations, *H. himera* also stood out as having the fewest facets (Fig. 1E).

Because elevation correlates with changes in climate and habitat type (25, 35), we further explored how facet count was influenced by the local environment. We modeled the relationships among forest cover and mean annual temperature, precipitation, and solar radiation and evaluated their effects on facet count independently of elevation (all four variables were negatively correlated with elevation; Fig. S4). Our structural equation model was well-specified (Fisher’s C = 4.28, df = 6, p = 0.64) and showed that facet count increased with mean annual temperature (p <0.001, Fig. S5A) and precipitation (p = 0.043; Fig. S5B), whereas solar radiation and forest cover, a measure of habitat continuity, had no significant effects (p <0.2; Figs. S5C, D). Together, these results show that variation in *H. erato* eye size is largely due to differences in facet count, and that higher elevations are associated with fewer facets. Elevation is likely a composite variable representing several environmental factors (Fig. S4), but temperature and precipitation were identified as important variables influencing eye morphology.

### Heritable changes in the visual system of *H. himera* are likely driven by selection

Having established population differences in eye morphology across the *erato* group, most notably in the high-elevation specialist *H. himera*, we next examined whether this variation is heritable, as previously reported for neural investment (33). Because population-level differences in sensory phenotypes are likely influenced by both genetic and environmental effects, we reared *H. himera* alongside *H. erato* from lower elevations under common-garden conditions in Ecuador. To further rule out that the differences we observe are not due to shared evolutionary history, we included both *H. erato cyrbia* (western clade) and *H. erato lativitta* (eastern clade) in our analyses (Table S2). Importantly, *H. himera* (eastern clade) is more closely related to *H. erato lativitta* than to *H. erato cyrbia* (Fig. 1B; (34)), though it meets both along narrow contact zones (36), and *H. erato lativitta* and *H. erato cyrbia* live in similar habitats (Figs. 2A, B).

**Figure 2.**
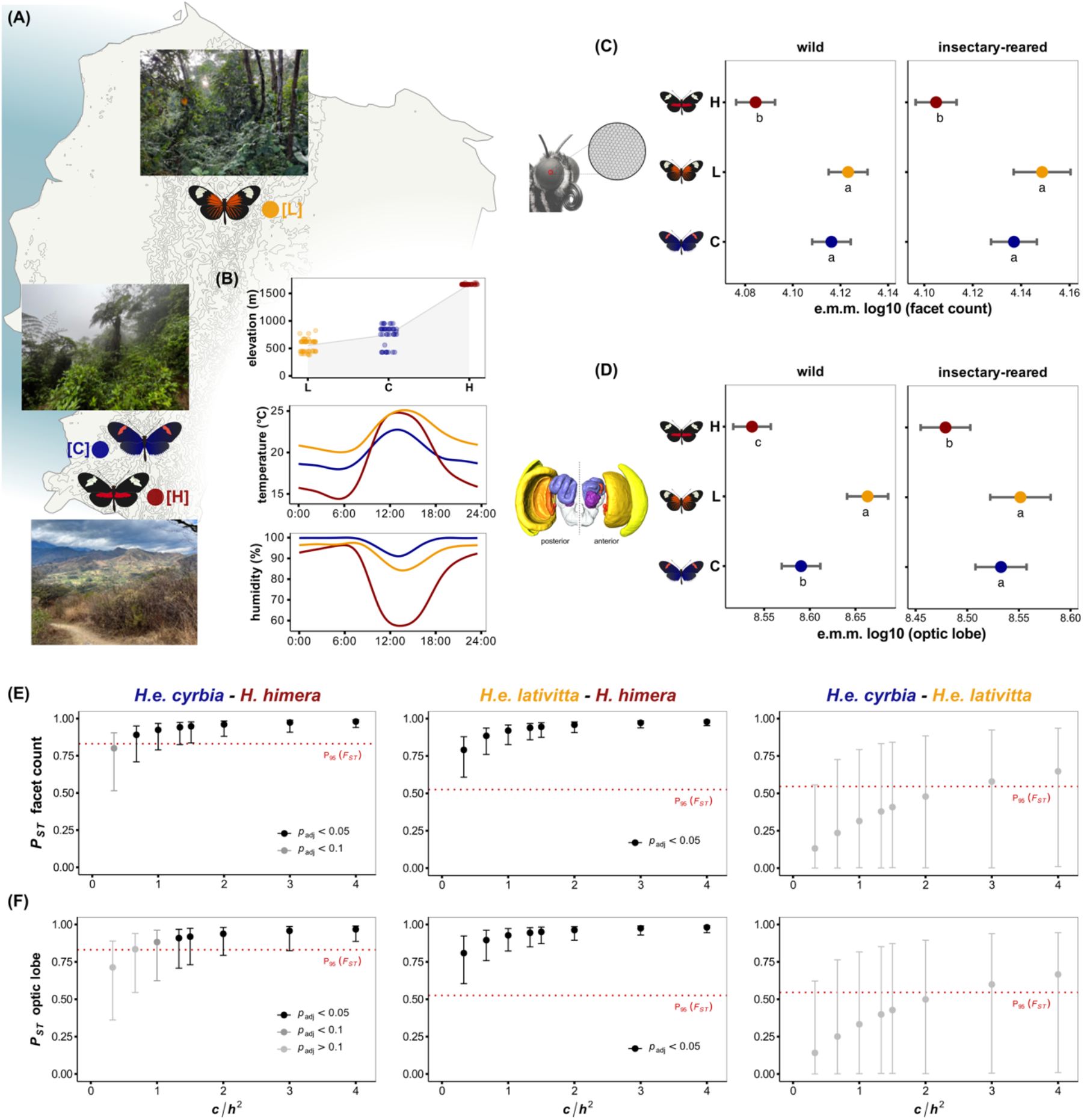
Visual system morphology is heritable and due to selection. **(A)** Source locations for the insectary populations of *H. erato cyrbia* [C], *H. erato lativitta* [L] and *H. himera* [H]. **(B)** Environmental conditions experienced by *H. erato cyrbia* and *H. erato lativitta* are similar, whereas *H. himera* lives at higher elevation and experiences more dramatic temperature fluctuations and lower humidity (see also pictures in A). Both wild and insectary-reared *H. himera* had reduced **(C)** facet count and **(D)** optic lobe volume compared to *H. erato cyrbia* and *H. erato lativitta*, suggesting trait heritability. Brain data for wild *H. erato cyrbia* and *H. himera* from (33). Estimated marginal means (e.m.m.) account for sex and an allometric control (tibia length for eyes, central brain volume for brains). Different letters indicate significant differences (Bonferroni-adjusted p <0.05), and the error bars represent 95% confidence intervals. *P_ST_*-*F_ST_* comparisons for **(E)** facet count and **(F)** optic lobe volume showed consistent evidence for divergent selection (*P_ST_* above the 95^th^ percentile of the *F_ST_* distribution [P_95_; red dotted line]) for *H. erato lativitta* vs *H. himera* and in most quantitative genetic scenarios (*c/h^2^* values) for *H. erato cyrbia* vs *H. himera*. Error bars represent 95% confidence intervals, and FDR-corrected p-values (p_adj_) calculated as the proportion of the *F_ST_* distribution above each *P_ST_* value. See also tables S9-S14.

Mirroring our results for wild-caught individuals (Fig. 2C; Table S3), insectary-reared *H. erato cyrbia* and *H. erato lativitta* had more facets than *H. himera* (p_adj_ <0.001) but did not differ from one another (p_adj_ = 0.23; Fig. 2C; Table S6). We also compared volumetric measurements of the optic lobe and its constituent neuropils (the major visual processing region of the insect brain; Fig. S6) for the three populations. Wild *H. erato lativitta* had larger optic lobes than both *H. himera* and *H. erato cyrbia* (p_adj_ <0.001), and *H. erato cyrbia* had larger optic lobes than *H. himera* (p_adj_ = 0.001; Fig. 2D; Table S4). Individual neuropil comparisons revealed more subtle variation (Table S5); the lamina and ventral lobula did not differ between *H. erato cyrbia* and *H. himera*, nor did the anterior optic tubercule - a central brain visual neuropil (p = 1 for all). In the insectary-reared samples, optic lobe volume did not differ between *H. erato cyrbia* and *H. erato lativitta* (p_adj_ = 0.6), but both had larger optic lobes than *H. himera* (p_adj_ <0.0013; Fig. 2D; Table S7) due to specific effects of the lamina, medulla, and lobula (Table S8). Finally, we observed a significant *population x sex* interaction (p = 0.011): for *H. erato cyrbia*, females had smaller optic lobes than males (p_adj_ <0.001), but the sexes did not differ for *H. erato lativitta* and *H. himera* (p_adj_ >0.4). Once again, differences in optic lobe volume were due to increases in the lamina, medulla, and lobula (Table S8).

To test for selection acting across the visual system, we calculated estimates of *P_ST_* for eye and brain morphology in our insectary populations. *P_ST_* is a phenotypic analogue of *F_ST_* that measures variation among populations relative to total variance across populations (37). Comparisons between *P_ST_* and *F_ST_* can be used as a test of divergent selection, where *P_ST_* values that exceed genome-wide *F_ST_* suggest greater phenotypic divergence than expected by neutral genetic divergence (37). Inferences made from *P_ST_* can be vulnerable to underlying assumptions regarding trait heritability (*h^2^*) and the proportion of the total variance that is presumed to be due to additive genetic effects across populations (*c*) (37). Heritability estimates for eye and brain morphology are unknown for *Heliconius*, so we used a wide range of *c/h^2^* ratios to test different quantitative genetic scenarios (see methods for complete description). After accounting for allometric effects, facet count *P_ST_* exceeded the 95^th^ percentile of genome-wide *F_ST_* in all but one scenario for *H. erato cyrbia* vs *H. himera* (Fig. 2E; Table S10), and comparisons with the individual neuropils indicate that the signal of selection is strongest for the medulla and, to a lesser extent, the lobula (Table S13). Optic lobe *P_ST_* also exceeded the 95^th^ percentile of genome-wide *F_ST_* under most quantitative genetic scenarios (potentially because of the medulla’s large effect on total optic lobe volume), though significance varied after correcting for multiple testing (Fig. 2F; Table S13). For *H. erato lativitta* vs *H. himera*, facet count *P_ST_* exceeded genome-wide *F_ST_* in every scenario (Fig. 2E; Table S11), and there was consistently strong evidence for selection driving variation in the medulla, lobula, and lobula plate, and to a lesser extent, the lamina and ventral lobe of the lobula (Table S14). Optic lobe *P_ST_* consistently exceeded the 95^th^ percentile of genome-wide *F_ST_* (Fig. 2F). Taken together, our results suggest an adaptive shift in the visual system of the high-elevation, dry-forest specialist *H. himera* compared to lower elevation *H. erato lativitta* and *H. erato cyrbia*.

### Components of the visual system can evolve independently

The second high-elevation specialist included in our wild survey, *H. chestertonii*, also tended to have fewer facets than the lower-elevation *eratos* (Fig. S1), though this was far less pronounced than for *H. himera* (perhaps due to a less extreme shift in habitat (29)). Previous results showed an adaptive shift in visual neuropil investment between *H. chestertonii* and its lower elevation relative, *H. erato venus* (Fig. 3C), with which it shares a contact zone in western Colombia (29, 32). These shifts mirror those in *H. himera* suggesting a degree of parallel evolution in neuropil investment (33). In contrast, we observed no significant differences in facet count for wild or insectary-reared *H. chestertonii* and *H. erato venus* (p >0.2, Figs. 3C, S7; see supplementary information for analyses of weak [p >0.05] *population x sex* interactions; Tables S18-S19). However, for both wild and insectary-reared butterflies (Figs. 3C, S7), *H. erato venus* had significantly larger corneal area than *H. chestertonii* (p <0.001; Tables S18-S19). This seems to reflect a broader shift in *H. erato venus*, in which the relationship between cornea area and facet count differs from the other sampled populations (Tables S3, S6, S15). Specifically, we observed a grade shift (α) in the relationship between corneal area and facet count for both wild (ANCOVA: F_1,308_ = 60.56, p <0.001; Fig. S8A) and insectary-reared *H. erato venus* vs the other *eratos* (ANCOVA: F_1,130_ = 129.19, p <0.001; Fig. S8B; Table S21), suggesting that *H. erato venus* has evolved an increase in mean facet size.

**Figure 3.**
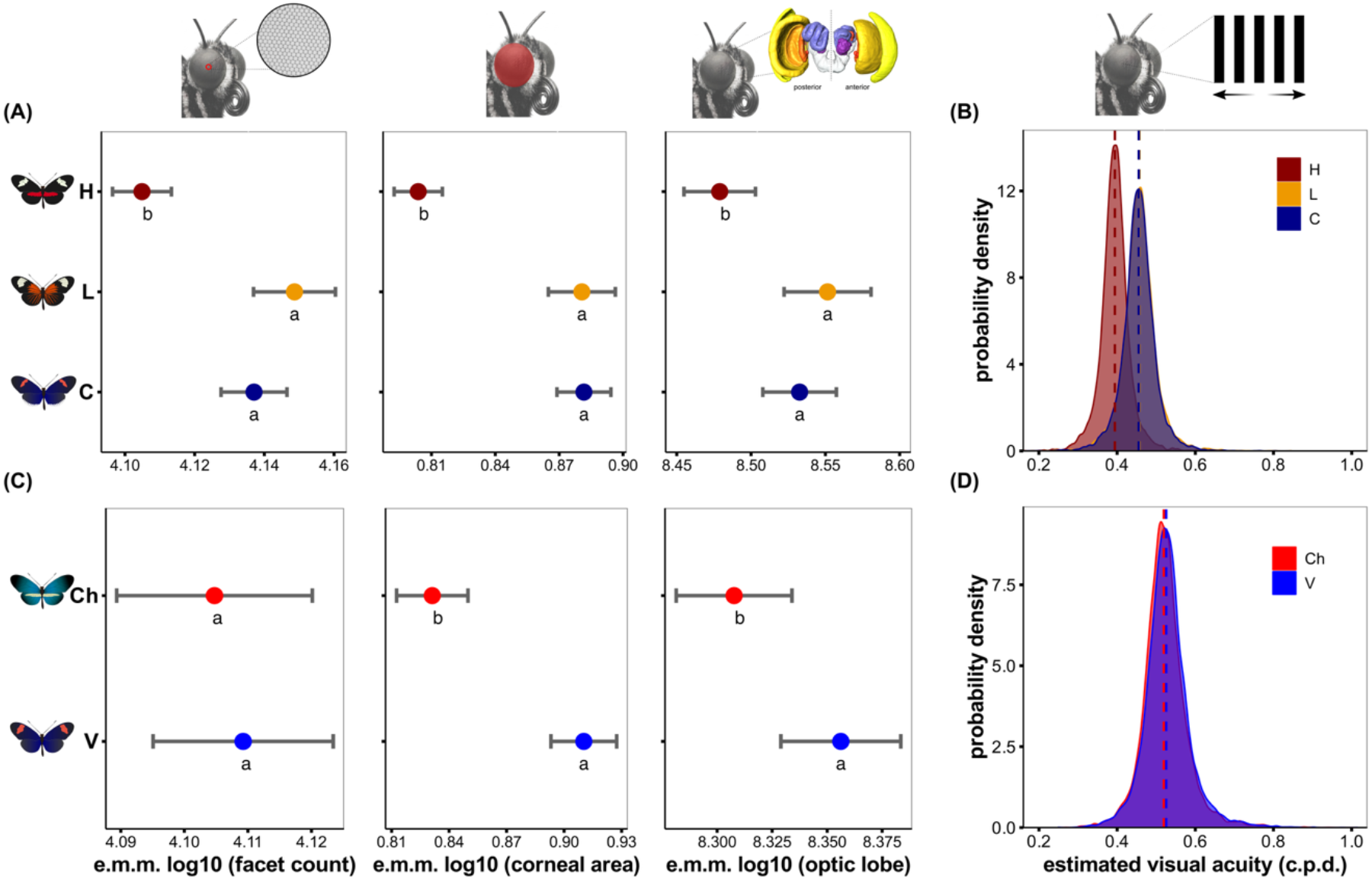
Visual traits can evolve independently. In Ecuador, **(A)** insectary-reared *H. himera* [H] (high elevation) had fewer facets, smaller corneal area, and smaller optic lobe volume than *H. erato cyrbia* [C] and *H. erato lativitta* [L] (both low elevation). **(B)** *H. himera* also had lower visual acuity. **(C)** In Colombia, insectary-reared *H. chestertonii* [Ch] (high elevation) had smaller corneal area and smaller optic lobes than *H. erato venus* [V] (low elevation), but facet count did not differ. **(D)** Visual acuity did not differ between *H. chestertonii* and *H. erato venus*. Estimated marginal means (e.m.m.) account for sex and allometry (tibia length or central brain volume), error bars represent 95% confidence intervals, and different letters indicate significant differences (Bonferroni-adjusted p <0.05). Distributions in B and D represent the posterior probability densities from Bayesian censored models, and the dashed vertical lines indicate the posterior medians (back transformed from log10 to the original scale, c.p.d.), accounting for sex and different observers. Brain data for insectary-reared *H. chestertonii* and *H. erato venus* in C from (32).

### Elevation-associated visual system divergence extends to spectral sensitivity

To complement our analyses of eye and brain morphology, we measured eyeshine in our five insectary populations. In the eyes of many butterfly species, lateral filtering pigments absorb short wavelengths and shift spectral sensitivity toward longer wavelengths, enabling long-wavelength sensitivity without functional changes to the opsins (17, 38–40). Eyeshine, created by unabsorbed light reflecting from the tapetum at the proximal end of each rhabdom (38), is an easily quantifiable proxy for lateral filtering pigments—ommatidia containing filtering pigments appear red (Fig. 4A). In Ecuador, we observed population-level differences in eyeshine: *H. himera* had fewer red-reflecting ommatidia than both *H. erato cyrbia* and *H. erato lativitta* in the frontal and ventral eye regions (p_adj_ <0.0035), whereas dorsal eyeshine did not differ (p = 0.9; Fig. S9B). Males had slightly more red ommatidia in the frontal eye region (p = 0.087), and females had significantly more red ommatidia in the ventral eye region (p = 0.014). In Colombia, *H. chestertonii* had fewer red ommatidia than *H. erato venus* in the frontal and ventral eye regions (p <0.001; dorsal eyeshine did not differ, p = 0.9), and there were no overall differences between the sexes (p >0.2; see supplementary results for *population x sex* interactions in both locations).

**Figure 4.**
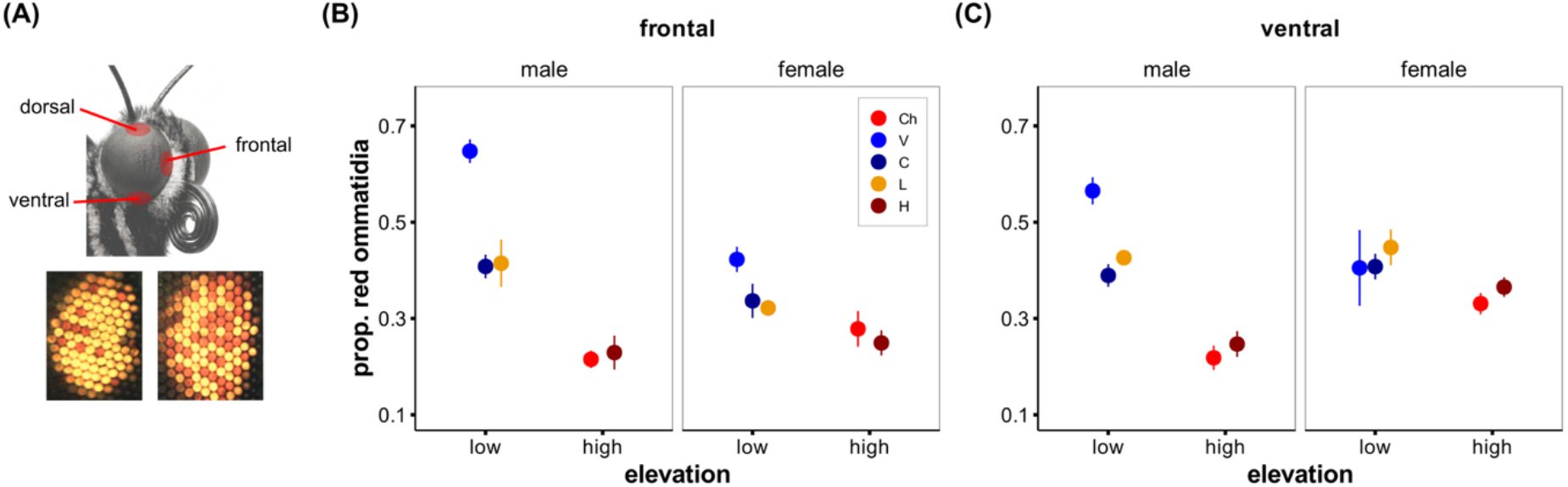
Spectral sensitivity represents another elevation-associated adaptation. Eyeshine, a proxy for lateral filtering pigments, across **(A)** different eye regions for insectary-reared butterflies from Ecuador (*H. himera* [H], *H. erato lativitta* [L], *H. erato cyrbia* [C]) and Colombia (*H. chestertonii* [Ch] and *H. erato venus* [V]). For both the **(B)** frontal and **(C)** ventral eye regions, high-elevation populations had fewer red-reflecting ommatidia than the low-elevation populations. Dorsal eyeshine did not differ. Points and error bars represent mean ± standard error. See supplementary results for expanded analyses within each location.

To test for shared patterns of ecologically driven divergence in lateral filtering pigments (following (32, 41)), we combined the data from Ecuador and Colombia and categorized elevation as low (*H. erato cyrbia*, *H. erato lativitta*, *H. erato venus*) or high (*H. chestertonii*, *H. himera*). We then tested how *elevation* influenced eyeshine while including *location* (Ecuador or Colombia) and *sex* as covariates. For the frontal and ventral eye regions, high elevation populations had significantly fewer red ommatidia (p <0.001; Fig. 4), and for the dorsal eye region, high elevation populations had slightly fewer red ommatidia (p = 0.091). Together, these results point to reduced lateral filtering pigments as a parallel adaptation to high-elevation habitats that is further modulated by regional (4) and sex-specific specialization across the eye.

### Peripheral and neural structures jointly determine visual acuity

The population-level discordance we observe between different visual traits presented an opportunity to explore how visual system differences may influence perception. To do this we used an optomotor assay to measure visual acuity across our five insectary populations (42). The optomotor response was elicited by rotating alternating vertical black and white stripes around a butterfly, repeated with increasingly narrower stripes until the butterfly failed to respond (i.e. the visual acuity threshold). This allowed us to compare individuals reared under common garden conditions in Ecuador or Colombia and see whether between population variation in eye and brain morphology corresponds to differences in visual acuity. As butterflies were tested to a maximum of 1.0 cycles per degree (c.p.d.), we used Bayesian censored regressions to analyze the behavioral data (see methods for details). In Ecuador, posterior estimates indicated that *H. himera* had lower visual acuity than both *H. erato cyrbia* (β = −0.062, 95% CI: [-0.110, −0.015]) and *H. erato lativitta* (β = - 0.064, 95% CI: [-0.112, −0.016]), whereas *H. erato cyrbia* and *H. erato lativitta* did not differ (β = 0.002, 95% CI: [-0.048, 0.050]; Fig. 3B). In Colombia, *H. erato venus* and *H. chestertonii,* which differed in cornea area and optic lobe investment, but not facet number, did not differ in visual acuity (β = 0.00, 95% CI: [-0.04, 0.05]; Fig. 3D).

To further assess how variation across specific visual phenotypes influences visual acuity, we generated hybrids between *H. erato cyrbia* and *H. himera* (Fig. 5A). This allowed us to experimentally dissociate individual components of the visual system from other species-specific traits. Using the same Bayesian censored regression framework, we first modeled the relationship between facet count and acuity across all available second-generation (F2) hybrids (n=186). Here, facet count showed a clear positive effect on visual acuity (β = 0.87; 95% CI: [0.15, 1.60]; Fig. 5C yellow bar, Fig. 5D). For a sub-sample of F2 hybrids for which we also had volumetric brain data (n=131; Fig. 5C, grey bars), we implemented a model evaluating the independent and interactive effects of facet count and optic lobe volume (after accounting for central brain volume and family identity, optic lobe volume was positively correlated with facet count: p <0.001; semi-partial *R^2^* = 0.136; Fig. S11A). Under our conservative prior framework [*b ∼ Normal(0, 0.5)*], the conditional main effects of facet count (β = 0.05; 95% CI: [-0.89, 0.98]) and optic lobe volume (β = −0.05; 95% CI: [-0.84, 0.73]) both overlapped zero. Because these parameters represent baseline slopes evaluated at zero on the log_10_ scale—a region outside our biological data range—these estimates safely default to the conservative prior. Crucially, the data shifted the posterior distribution away from the prior for the interaction term, revealing a positive *facet count x optic lobe* interaction (β = 0.18; 95% CI: [0.02, 0.33]; Fig. 5C). This positive interaction demonstrates a mutual dependence between eye and brain morphology, where the positive relationship between either trait and visual acuity is contingent upon the concurrent expansion of the other (Fig. 5E).

**Figure 5.**
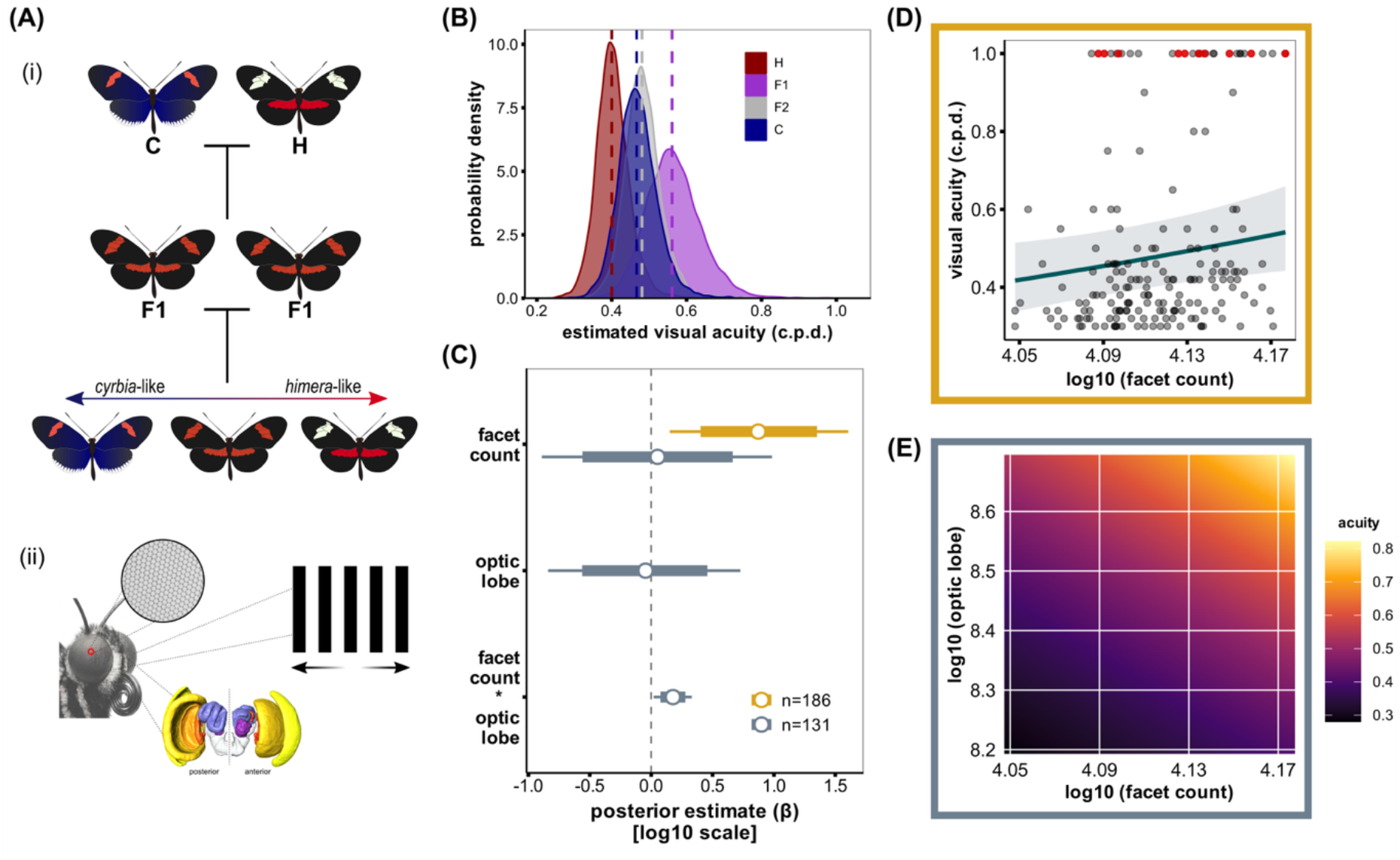
Facet count and neuroanatomy jointly influence visual acuity. **(A)** We generated hybrids of *H. erato cyrbia* [C] and *H. himera* [H] and assessed each for eye and brain morphology and **(B)** visual acuity (see supplementary results for pairwise comparisons). The shaded distributions represent the posterior probability densities from the Bayesian censored model, and the dashed vertical lines indicate the posterior medians (back transformed from log10 to the original scale, c.p.d.). **(C)** For 186 F2 hybrids, facet count positively influenced visual acuity (yellow bar), whereas a smaller sub-sample of F2s with complete brain data (n=131, grey bars) revealed an interactive effect of facet count and optic lobe volume. Horizontal bars represent the 80% (thick) and 95% (thin) posterior intervals, with the central point indicating the posterior median. **(D)** The positive association between facet count and visual acuity for the full dataset (n = 186). The solid line and shaded ribbon represent the back-transformed posterior mean and 95% credible interval (see also yellow bar in C), and the points are raw observations [red points represent F2 hybrids that were tested >1.0 c.p.d.; see methods]. (**E)** The interactive effect of facet count and optic lobe on visual acuity in the smaller sub-sample of F2s (n=131; see also grey bar in C).

Analyses with the individual neuropils suggest that this effect of the optic lobe may be driven by the lamina (*facet count x lamina* interaction: β = 0.15; 95% CI: [- 0.01, 0.31]), which was more strongly correlated with facet count than total optic lobe volume (after accounting for central brain volume and family identity: p <0.001; semi-partial *R^2^* = 0.274; Fig. S11B). We also observed slight effects of the accessory medulla (β = 0.23; 95% CI: [-0.07, 0.53]), but it was not correlated with facet count (p = 0.4) and its direct role in visual perception is unlikely ((43, 44); see also the supplementary results, and discussion below). Collectively, our isolated analysis of the larger dataset confirms that facet count is a major driver of visual acuity, and our integrative models show that this structural investment is computationally limited, requiring a corresponding expansion in downstream neural architecture.

## Discussion

While natural selection is expected to shape visual systems, how distinct components of the visual pathway co-vary to drive divergence across terrestrial populations has remained largely untested. By leveraging continental-scale sampling and our ability to raise and cross ecologically divergent *Heliconius* populations, our results reveal heritable variation both within and between species across multiple components within the visual system. Our population comparisons show that these different components can evolve independently, but together are broadly associated with elevation, representing a major axis of ecological transition. Finally, both eye and brain morphology, independent of body size, shape visual acuity, likely influencing visual perception across diverse forest types.

Elevation represents a major ecological transition, especially in the Andean tropics (25, 35), influencing the visual environment in which animals must perform key behaviors. Across *Heliconius erato* populations, we saw a striking reduction in facet count with increasing elevation. This negative correlation aligns with a growing body of evidence documenting altitudinal adaptations across this genus (26–29, 31), including reduced visual neuropil investment in the high-elevation specialists *H. himera* and *H. chestertonii* (32, 33). Crucially, the reduction in peripheral visual investment we observe cannot be explained by allometric scaling (Fig. S3), but instead reflects a targeted, adaptive down-scaling of the visual system at higher elevations. Although elevation was a significant predictor of facet count, we expect that elevation *per se* is not the causal factor influencing peripheral morphology. Instead, elevation likely acts as a proxy for habitat-type, with concurrent changes in several environmental factors that together influence the local forest habitat (Fig. S4). The significant effects of temperature and precipitation suggest that other micro-climatic or biotic variables not captured in our model—such as plant density, vegetation type, or the ambient light spectra—may be the proximate drivers of facet count divergence (Fig. S5).

We also provide evidence for elevation-associated variation in lateral filtering pigments: both high-elevation residents, *H. himera* and *H. chestertonii*, possessed fewer red ommatidia than the lower-elevation populations. Within the *erato* species complex studied here, this variation may reflect adaptation to the local light conditions. Compared to open habitats, forested environments possess proportionally more long-wavelength light (6), and the greater abundance of lateral filtering pigments among lower-elevation *H. erato* may tune spectral sensitivity to the wavelengths most prevalent in forests. Increased long-wavelength sensitivity may also amplify color contrast against a complex forest background (45). However, this is somewhat difficult to reconcile with results from more distantly related *Heliconius* (17), or other Nymphalids (5), where species in more open habitats have more red-reflecting lateral filtering pigments. Possibly, increased filtering pigments allows better color discrimination in *Heliconius*, which may be beneficial for males that rely on colour to find suitable mates. This could be particularly valuable within more diverse lower-elevation, forest-edge communities (10), inhabited by both *H. erato* and *H. melpomene*, where males must discriminate among females of multiple co-occurring *Heliconius* species. Regardless, these contrasting patterns suggest that the adaptive value of filtering pigments may vary among lineages, even within *Heliconius*, depending on their interaction with species-specific ecology and behavior.

The high-elevation specialist *H. himera* stood out for its reduced facet count and reduced investment in several optic neuropils in comparisons involving both wild and insectary-reared butterflies. These heritable differences seem to represent whole visual system adaptation. This is supported by our tests of selection, where *P_ST_* exceeded genome-wide *F_ST_* across a broad range of quantitative genetic scenarios, and previous results implicating selection during *Heliconius* eye (20) and brain evolution (23, 32). However, our results also suggest that the peripheral eye morphology and visual neuropils can evolve as independent, decoupled traits. Although shifts in neuroanatomy have been identified between high-elevation *H. chestertonii* and low-elevation *H. erato venus* from Colombia (32), we did not observe any corresponding shift in facet count. Notably, *H. erato venus* has significantly larger corneal areas, as well as a pronounced grade shift in the relationship between corneal area and facet count, relative to all other populations tested (Fig. S8). While we did not measure individual facet diameters, this strongly implies a larger mean facet size for *H. erato venus*, perhaps reflecting an alternative sensory strategy. Unlike the other populations included in our study, *H. erato venus* was sampled from coastal habitats. This environment potentially introduces several visual challenges—such as high-intensity surface glare from water and sand, alongside abrupt spatial transitions between deep forest shade and open shorelines—that are largely absent in interior tropical rainforests. These conditions may favor larger facets to optimize light capture and, together with higher lateral filtering abundance, may help maximize the visual signal-to-noise ratio (46) in this coastal environment. Thus, while clear evolutionary patterns exist within the visual system, our results highlight how localized ecological pressures can drive complex and divergent sensory strategies. Future sampling of other coastal *Heliconius* populations and/or *H. erato venus* from additional areas (24) will help to explore these patterns further.

We confirmed the behavioral consequences of these morphological shifts using an optomotor assay to assess visual acuity. In Ecuador, *H. himera* exhibited lower visual acuity than both *H. erato cyrbia* and *H. erato lativitta*, matching its lower facet count. In Colombia, the visual acuities of *H. chestertonii* and *H. erato venus* did not differ, matching their equivalent number of facets (Fig. 3). These behavioral patterns suggest that visual acuity may be more constrained by peripheral facet number than neural investment (i.e. both high-elevation specialists possess reduced visual neuropils, but lower acuity was observed only for *H. himera* with fewer facets). Possibly, this is because the peripheral optics represent the first limiting step in the visual pathway (47), and visual behavior is largely governed by the quantity and quality of information captured by the external receptors.

To directly test the relationship between visual acuity and visual system morphology, we evaluated second-generation hybrids between *H. erato cyrbia* and *H. himera*, thereby experimentally separating different segregating components of the visual systems. Facet count positively predicted visual acuity, consistent with physiological models of compound eyes (48), and representing the first large-scale experimental validation of this relationship in insects. Furthermore, volumetric brain data from a sub-sample of hybrids revealed that visual acuity is modulated by a positive interaction between facet count and optic lobe volume. Finer-scale analysis with the individual neuropils confirmed a similar, albeit slightly weaker interaction between facet count and the lamina (see supplementary results), the first optic neuropil to receive photoreceptor input and a primary processor for motion vision. In *Drosophila*, responsiveness to moving gratings has been attributed to lamina monopolar cells (LMC) L1 and L2 (49), and in Lepidoptera, L1/L2 are similarly involved in motion vision (50, 51). Our analyses also suggest that the accessory medulla might play a role, but this neuropil governs circadian control in insects (43) and does not directly respond to moving stimuli (44). Thus, the accessory medulla’s impact on acuity is likely indirect, potentially due to its role in gating daily visual sensitivity thresholds. Nonetheless, the stronger interaction effect observed for facet count and total optic lobe volume, rather than the individual neuropils, implies that multiple brain regions contribute to variation in visual acuity. Ultimately, our results show that variation in facet count directly affects visual acuity, but this peripheral investment does not function in isolation: improving acuity across evolutionary time fundamentally requires matching variation in downstream neural infrastructure.

## Methods

### Wild butterflies

We collated samples of wild *Heliconius erato* spp. (see Table S1 for sample overview). For all samples, whole butterfly bodies were preserved in DMSO/EDTA/NaCl and frozen, and all relevant metrics were recorded (e.g. identity, sex, GPS coordinates of capture).

### Insectary-reared butterflies

#### Ecuador

At the Universidad Regional Amazónica Ikiam in Tena, Ecuador (0°56’53.7"S, 77°51’56.0"W), we established outbred stock populations of *H. erato cyrbia*, *H. erato lativitta*, and *H. himera* from wild-caught butterflies. Both *H. erato cyrbia* and *H. himera* were collected in Southern Ecuador, near Balsas (3°43’26.0"S, 79°50’52.1"W) or Buenaventura (3°39’06.7"S, 79°45’56.3"W) and Vilcabamba (4°15’13.921’’S, 79°12’ 41.461’’W), respectively. *H. erato lativitta* was collected locally around Tena. All butterflies were maintained under common garden conditions in outdoor insectaries in 2 x 2 x 2.3 m cages, where they were provided with 20% sugar solution and *Lantana* sp. and *Psiguria* sp. flowers. Eggs were collected regularly from *Passiflora punctata* hostplants, the larvae were individually reared in pots with fresh hostplant leaves, and all butterflies were marked with unique identification codes on their wings after eclosion.

We also generated hybrids of *H. erato cyrbia* and *H. himera*. First-generation (F1) hybrids were generated from fourteen crosses of a *H. erato cyrbia* female x *H. himera* male and five crosses of a *H. himera* female x *H. erato cyrbia* male. Second-generation (F2) hybrids were created by mating F1s (24 crosses). All hybrids were reared under the same insectary conditions as the parental species.

#### Colombia

At the José Celestino Mutis Experimental Field Station (Universidad del Rosario), near La Vega, Colombia (5.0005° N, 74.3394° W), we established outbred stock populations of *H. erato venus* [collected from Buenaventura (03°50’0.04” N, 77°15’45.1” W)] and *H. chestertonii* [collected from Montañitas (3°40’36.3" N, 76°31’19.4" W)]. Butterflies were reared under common garden conditions, in outdoor insectaries (1 x 3 x 2 m cages) with access to 20% sugar solution and *Gurania*, *Lantana* and *Psiguria* spp. flowers. Eggs were collected from *Passiflora biflora* hostplants, individual larvae reared in pots, and all butterflies were marked with unique identification codes on their wings.

### Eye morphology

Quantification of eye morphology followed previously established protocols (20). In brief, frozen specimens were thawed, and the eyes and hind legs were dissected out. The legs were immediately imaged, while the eyes were placed in 20% NaOH overnight to loosen the tissues behind the cuticular cornea. The following day, the eye cuticle was cleaned of excess tissue and mounted on a microscope slide. After drying, the cuticle was imaged, and facet count, corneal area, and tibia length were measured in ImageJ (52). Full protocols are available online (53, 54).

### Brain morphology

Complete heads were removed from freshly sampled butterflies, fixed in zinc-formaldehyde solution (ZnFA; 0.25% ZnCl2, 0.788% NaCl, 1.2% sucrose, 1% formaldehyde) for 16–20 hours at room temperature under gentle agitation, and then incubated in 80% methanol / 20% DMSO for 2 hours. Samples were subsequently transferred to 100% methanol and stored at −20 °C until processing.

Brain morphology was visualized using immunofluorescence staining following established protocols (55, 56). Brains were rehydrated through a graded methanol series (90%, 70%, 50%, 30%, and 0%, each in 0.1 M Tris buffer, pH 7.4; 10 minutes per step). Tissues were pre-incubated for 2 hours at room temperature in 5% normal goat serum (NGS) diluted in PBS containing 1% DMSO and 0.005% sodium azide (PBSd-NGS). Primary antibody 3C11 (anti-synapsin, Developmental Studies Hybridoma Bank, RRID: AB_2315424) was applied at a 1:30 dilution in PBSd-NGS and incubated for 3.5 days at 4 °C under continuous agitation. Following three 2-hour rinses in PBSd, samples were incubated with a Cy3-conjugated goat anti-mouse IgG secondary antibody (Jackson ImmunoResearch; Cat. No. 115-165-146; RRID: AB_2338690), diluted 1:100 in PBSd-NGS, for 2.5 days at 4 °C under agitation. After staining, samples were dehydrated through an ascending glycerol series (1%, 2%, 4% for 2 hours each, followed by 8%,15%, 30%, 50%, 60%, 70%, and 80% for 1 hour each) in 0.1 M Tris buffer with 1% DMSO, then transferred to 100% ethanol and cleared in methyl salicylate.

Confocal imaging was performed using a Stellaris 5 confocal laser-scanning microscope (Leica Microsystems, Mannheim, Germany) with a 10× dry objective (NA 0.4; Leica Material No. 11506511), a mechanical z-step of 2 μm, and an x–y resolution of 512 × 512 pixels. Whole-brain stacks were acquired as image stacks (20% overlap) which were automatically merged. The z-dimension was scaled by a factor of 1.52 to correct for artifactual compression associated with the use of a 10× air objective (55).

We measured the volumes of optic and central brain neuropils, including the lamina (LAM), medulla (ME), lobula (LOB), lobula plate (LOP), accessory medulla (aME), ventral lobula (vLOB), antennal lobe (AL), anterior optic tubercle (AOTU), and mushroom bodies (MB). The rest-of-central-brain (rCBR), calculated by subtracting the above segmented neuropils from the total central brain volume, was used as an allometric control. Paired structures were quantified in one hemisphere and doubled. Brain regions were segmented from confocal image stacks using Amira v.2023.2 (Thermo Fisher Scientific). To accelerate segmentation, we applied convolutional neural networks using Biomedisa (57) to semi-automatically delineate neuropil boundaries. The trained networks [presented in full in (21)] were used to generate initial segmentations, which were manually refined in Amira to ensure anatomical accuracy.

### Environmental data

For each wild butterfly, we confirmed elevation from the recorded GPS coordinates of capture using the OpenTopoData API (https://api.opentopodata.org/), querying the NASA Shuttle Radar Topography Mission (SRTM) 30m (1 arc-second) global dataset (58). For sampling locations of *H. erato cyrbia* (Balsas), *H. erato lativitta* (Tena), and *H. himera* (Vilcabamba) in Ecuador, we also tracked temperature and humidity for ∼one year (March 2022 - March 2023) via HOBO Pro v2 temp/RH loggers (Onset Computer Corporation) set to measure temperature and relative humidity every 30 minutes. The data is plotted as the mean 24-hour variation over the one-year period (Fig. 2B).

### Tests of selection

As in prior work (20), we calculated *P_ST_* values using *pst()* (Pstat package (59)) and used hind tibia length for allometric corrections [*res()*]. *P_ST_* calculations include the ratio *c/h^2^*, where *c* represents the proportion of the total variance due to additive genetic effects across populations and *h^2^* is heritability (37). Heritability estimates for *Heliconius* eye and brain morphology are unknown, so in addition to the default value of 1 (common for morphological traits (37)), we used *c/h^2^* ratios ranging from 0.33 to 4 (as in (20, 23)).

The distribution of windowed *F_ST_* values at 4-fold degenerate sites derives from genome-wide polymorphism data including wild individuals of *H. erato cyrbia* (n = 15), *H. erato lativitta* (n = 27), and *H. himera* (n = 31) (Table S9). Illumina short reads were processed with *bbduk* from BBMap/BBTools (60) for quality and adaptor trimming and mapped to the *H. erato demophoon* reference genome (v1) (61; LepBase, downloaded 29 August 2023) using BWA-MEM (v0.7.17) (62). Only properly paired reads with a mapping quality (MAPQ) ≥20 were retained. Duplicated reads were subsequently removed using the *MarkDuplicates* function from Picard Tools (v2.0.1) (63). Variant calling was conducted using *Freebayes* (v1.0.2) (64). Biallelic SNPs and invariant sites were retained, and the remaining sites were filtered sites with a site quality >10, missingness <0.3, and mean depth of coverage between 10 and 50 using bcftools (v1.19) (65). The annotation of the *H. erato demophoon* reference genome (66) (https://zenodo.org/records/11519658) was processed with the python script *codingSiteTypes.py* (https://github.com/simonhmartin/genomics_general) to identify and keep only 4-fold degenerate sites for downstream analyses. Finally, estimates of *F_ST_* were obtained for non-overlapping 10 kb windows using *pixy* (1.2.7.beta1) (67).

We calculated p-values as the proportion of the *F_ST_* distribution that was above each *P_ST_* value; values above the 95^th^ percentile of the *F_ST_* distribution were interpreted as an indication of selection (20, 68). We also present the 95% confidence intervals of the *P_ST_* point estimates to account for uncertainty in *c* and *h^2^* (37).

### Visual acuity

We used an optomotor assay to measure behavioural visual acuity (42). The device consisted of interchangeable visual stimuli of alternating vertical black and white stripes that rotated around a fixed base. The width of one cycle (a set of alternating black and white stripes) was calculated as cycle width (mm) = [(C/360) / a], where ‘C’ is the circumference of the experimental arena and ‘a’ is the intended visual acuity in cycles-per-degree (c.p.d.) (69). For most butterflies (n = 563), we used stimuli with spatial frequencies of 0.3 c.p.d. (cycle width = 9.95 mm) to 1.0 c.p.d. (cycle width = 2.98 mm), though 16 hybrids were opportunistically tested to 1.6 c.p.d. (cycle width = 1.87 mm). For each trial, responsiveness was first confirmed at 0.3 c.p.d. As visual acuity depends on the distance between the stimulus and the perceiver, all butterflies were restrained in a clear PLEXIGLAS cylinder (4 cm radius; 15 cm height) at the center of the base. All assays were conducted inside, illuminated by an overhead LED ring lamp, and video recorded from above. A positive response was scored if the butterfly changed orientation of its head/antenna in the direction of the moving stimulus in two consecutive stimuli rotation reversals. All positive responses were confirmed from videos, and butterflies with unclear responses were excluded from the experiment.

### Eyeshine

Eyeshine measurements followed previously established protocols (4, 5). In short, butterflies, held in a slotted plastic tube and immobilized with plasticine, were suspended from the arm of a micromanipulator on a goniometric cradle, and the head was oriented towards a custom-built ophthalmoscope (4). Focus was adjusted on the cornea until the optical axes of several ommatidia were directly facing the objective lens (10×0.25 NA; Plan N, Olympus, Tokyo, Japan), thus viewing the pseudopupil. After dark adaptation, a broadband (470–850 nm) LED (MBB1F1 Thorlabs, Munich, Germany) was switched on to reveal eyeshine. The light was projected along an optical fiber, and a beam-splitter (Thorlabs, Newton, MA, USA) provided co-axial illumination to the eye and attached UI-3590CP-C-HQ-R2 camera with a CMOS colour sensor (Imaging Development Systems, Germany). Eyeshine was recorded with the uEye Cockpit software (IDS Software Suite 4.95) and quantified as the proportion of red ommatidia.

### Statistical analysis

#### Eye and brain morphology

We used linear mixed models [*lmer()*] in R to explore how wild *H. erato* eye morphology (facet count or corneal area) varied across populations, while accounting for sex, tibia length and sampling location (as a random effect). For comparisons within Ecuador and Colombia (wild and insectary-reared), we used linear models [*lm()*] to test how eye morphology or neuropil volume differed between populations, while accounting for sex and an allometric control (tibia length or rCBR). For analyses including *cyrbia-himera* hybrids, we also included family identify as a random effect in the *lmer()* models. In all comparisons, log10-transformations were used to normalize the residuals around the allometric relationships to meet the assumptions of normality (70). Model assumptions were confirmed via visual inspection (residual vs. fitted and normal Q-Q plots). The significance of fixed effect parameters was determined by likelihood ratio tests via *drop1()*, and minimum adequate models (MAM) were selected using statistical significance (71, 72). We used *Anova()* (car package (73)) to estimate significant fixed effect parameters, and in the case of more than two categories per fixed effect parameter, we conducted posthoc comparisons with *pairs()* (emmeans package (74)) and report p-values adjusted for multiple comparisons (adjust = “bonferroni”). To accurately visualize multiple significant fixed effects, we extracted and plotted the estimated marginal means from each MAM using the *emmeans* package (74).

We also explored whether the scaling relationships between facet count and corneal area differed for *H. erato venus* vs the other *eratos* (*H. erato venus* had larger corneal area than *H. chestertonii*, but facet count did not differ; see results). For both wild and insectary-reared butterflies (Colombia and Ecuador), we grouped butterflies as *H. erato venus* or ‘other’ and then tested the linear model: *log10 (corneal area) ∼ log10 (facet count) * group + log10 (tibia length)*. The *facet count x group* interaction was consistently non-significant (p >0.13), indicating similar slopes, and was dropped from the models. Significance for the independent effect ‘group’ was interpreted as evidence for a grade shift (α) in the relationship between facet count and corneal area.

To examine the relationships between eye and brain morphology, we built *lmer()* models accounting for central brain volume and family [e.g. *log10(facet count) ∼ log10(optic lobe) + log10(rCBR) + (1|familiy)*]. We estimated full model marginal *R^2^* and optic lobe- or lamina-specific semi-partial *R^2^* using *partR2()* (75)

#### Environmental effects

To evaluate the direct and indirect effects of the environment on facet count, we implemented a piecewise structural equation modeling framework [*psem()* (76, 77)]. For each sampled butterfly (GPS coordinates per individual), we incorporated mean annual temperature, precipitation, and solar radiation from WorldClim (30-arc sec resolution [∼1 km^2^] (78)) and forest cover (0.3 arc-second resolution [∼9m]) from ESA WorldCover via the *landcover()* function (79). This allowed us to model hierarchical relationships between the environmental variables (independent of elevation; all four variables are negatively correlated with elevation; Fig. S4) using a series of structured *lmer()* models. Our paths were: 1) *solar radiation ∼ temperature + precipitation + (1|population)* and 2) *forest cover ∼ temperature + precipitation + solar radiation + (1|population)*, followed by 3) *facet count ∼ temperature + precipitation + solar radiation + forest cover + sex + tibia length + (1|population)*. Finally, to account for minor sampling imbalances across habitat types, a correlated error relationship was specified for *sex ∼ forest cover*. To visualize and interpret the continuous partial effects of each variable on facet count, we calculated estimated marginal means (e.m.m.) using *ggpredict()*(77).

#### Visual acuity

We analyzed visual acuity differences between populations and sexes using Bayesian censored models (*brm()*, brms package (80)), with 4,000 iterations per chain, including a 1,000-iteration burn-in and no thinning (thin=1), for a total of 12,000 post-warmup draws. Most butterflies (n = 563) were tested to a maximum of 1.0 c.p.d., and we applied right censoring when butterflies achieved 1.0 c.p.d. Sixteen *cyrbia-himera* hybrids were opportunistically tested to a maximum of 1.6 c.p.d. (n_F1_ = 4; n_F2_ = 12), nine of which reached the highest threshold (n_F1_ = 3; n_F2_ = 6). For these butterflies, we applied right censoring when visual acuity = 1.6 c.p.d. Acuity was log10-transformed and *family = gaussian()*. We used weakly informative priors: regression coefficients (slopes) were assigned a *Normal(0, 0.5)* distribution and the intercept a *Normal(−0.36, 1)* distribution (centered on the mean log10(visual acuity) value). Residual noise (**σ**) was assigned a *Student-t(3, 0, 0.5)* prior. To account for different experimental observers (as a random effect), the group-level standard deviation (*sd*) was assigned a *Half-Normal(0, 0.1)* prior. Model performance was evaluated using *pp_check()* to ensure the model captured the distribution of the observed data, and brms::*pairs()* was used to assess convergence and sampling.

We used the same approach to test the relationship between visual acuity and eye/brain morphology in the F2 hybrids. The models included facet count or neuropil volume (or both), an allometric control (tibia length, rCBR, or both), and family identity as random effect. As above, all variables were log10-transformed and *family = gaussian()*. We used slightly modified priors: group-level (family identity) *sd* was assigned an *exponential*(*2*) prior. Sex and observer were not included in the F2 analyses as model comparison [*loo()*] showed both to be unnecessary (their inclusion did not increase predictive performance).

#### Lateral filtering pigments

Using the *lm()* approach detailed above, we tested how the proportion of red ommatidia in each eye region (frontal, ventral, dorsal) was influenced by population and sex.

To test for shared patterns of ecologically driven divergence in lateral filtering pigment abundance (as in (32, 41)), we combined data from Ecuador and Colombia and categorized *elevation* as low (*H. erato cyrbia, H. erato lativitta, H. erato venus*) vs high (*H. himera, H. chestertonii*). We tested [*lm()*] if variation in lateral filtering pigments was attributable to *elevation*, while controlling for differing evolutionary histories (*location*; Colombia or Ecuador) and *sex* (both variables were included as fixed effects).

## Acknowledgements

<u>Ecuador</u>: We thank Gladis Grefa and Whitney Wright for their contributions to plant and butterfly stock maintenance. Butterflies were collected with permission of the Ministerio del Ambiente, Agua y Transición Ecológica (MAATE-DBI-CM-2021-0176; MAATE-DNB-CM-2021-0176) and the Fundación de Conservación Jocotoco (for collecting within Buenaventura Reserve). <u>Colombia</u>: We thank Universidad del Rosario for insectary access at José Celestino Mutis Experimental Field Station and Isabel León for technical assistance. Field collections were conducted under permit no. 530 issued by the Autoridad Nacional de Licencias Ambientales (ANLA) of Colombia. <u>Germany</u>: Katerina Altouva, Florian Nebel, and Berkay Atalas assisted with sample processing. Microscopy was performed at the Center for Advanced Light Microscopy (CALM), LMU Munich. The Leica Stellaris 5 confocal microscope used in this study was funded by the German Research Foundation (Project number 495215303). Computing was performed on the BioHPC hosted at Leibniz Rechenzentrum Munich funded by German Research Foundation (INST 86/2050-1 FUGG). This research was supported by a European Research Council starting grant (grant no.: 851040) to RMM, a Natural Environment Research Council Independent Research Fellowship (NE/N014936/1) to SHM, and a Natural Environment Research Council GW4+ Doctoral Training Partnership studentship to JBW. Gemini 3 Flash (April 2026) was used to generate the butterfly head illustration used in figures 1-5.

## Data, Materials, and Software Availability

All raw data and analysis scripts are available on Zenodo (https://doi.org/10.5281/zenodo.22129140), and newly generated whole-genome sequences will be available on the European Nucleotide Archive (ENA; project accession: PRJEB124587).

## Author contributions

Conceptualization: DSW, SHM, RMM

Data curation: DSW, JB, YPT, LA, JG-V, LQ, EMP, MG-R, PC-V, DL-U, GR-M

Formal analysis: DSW, RMM

Funding acquisition: RMM, SHM

Investigation: DSW, JB, YPT, LA, AM, BW, JG-V, LQ, EMP, MG-R, PC-V, DL-U, GR-M

Methodology: DSW, JB, YPT, AM, BW, JG-V, SHM, RMM

Project administration: DSW, LQ, CP-D, CS, CNB, SHM, RMM

Resources: PS-C, NJN, CJ, CP-D, CS, CNB, SHM, RMM

Software: JB, JG-V

Supervision: DSW, SHM, RMM

Validation: DSW, RMM

Visualization: DSW, RMM

Writing – original draft: DSW, SHM, RMM

Writing – review & editing: all authors reviewed and edited

