## Supplementary information for "Adaptation in the eye and brain contributes to species divergence in visual perception in *Heliconius* butterflies"

### Supplementary results

**Corneal area, but not facet count, differed for *H. erato venus* vs *H. chestertonii*.** In Colombia, prior work documented smaller visual neuropils for high-elevation *H. chestertonii* vs low-elevation *H. erato venus* (1). In the present study, there were no population-level differences in facet count for wild or insectary-reared *H. chestertonii* and *H. erato venus* ($p > 0.2$; Fig. S7). For the wild butterflies, there was a weak *population x sex* interaction (LM: $F_{1,27} = 2.93$, $p = 0.098$), and posthoc comparisons showed slightly more facets for *H. erato venus* males than *H. chestertonii* males (estimate = 0.026, $p_{adj} = 0.056$; females did not differ, $p_{adj} = 0.5$; Fig. S7A). The same interaction could not be evaluated in the insectary as we had few female samples (1 *chestertonii*, 3 *venus*; Table S2). At the population level, wild *H. erato venus* had significantly larger corneal area than *H. chestertonii* (LM: $F_{1,28} = 18.11$, $p < 0.001$). Corneal area was also influenced by a slight *species x sex* interaction ($p = 0.064$); posthoc comparisons showed larger corneas for *H. erato venus* males compared to *H. chestertonii* males ($p_{adj} < 0.001$). Females did not differ ($p_{adj} = 0.17$) but trended in the same direction (Fig. S7B). Patterns were similar in the insectary; *H. erato venus* had larger corneal area than *H. chestertonii* (LM: $F_{1,30} = 56.78$, $p < 0.001$; Fig. S6D). This suggests variable facet size (larger diameter) for *H. erato venus* (see Fig S8).

**Elevation-associated visual system divergence extends to spectral sensitivity (continued).** In Ecuador, *H. himera* had fewer red-reflecting ommatidia than both *H. erato cyrbia* and *H. erato lativitta* in the frontal eye region ($p_{adj} < 0.0019$). Ventral eyeshine was influenced by a marginally non-significant *population x sex* interaction ($p = 0.089$); *H. himera* males had fewer red ommatidia than *H. erato cyrbia* and *H. erato lativitta* ($p_{adj} < 0.008$), whereas female eyeshine did not differ ($p_{adj} > 0.7$; though *H. himera* vs *H. erato lativitta* was marginally non-significant, $p_{adj} = 0.08$). There were no differences in dorsal eyeshine ($p = 0.9$; Fig. S9B). In Colombia, frontal eyeshine was influenced by a significant *population x sex* interaction (LM: $F_{1,20} = 20.71$, $p < 0.001$): *H. chestertonii* males had fewer red ommatidia ($p < 0.001$), and this difference was greater than in females (though *H. chestertonii* females also had fewer red ommatidia; $p = 0.005$). There was also a significant *population x sex* interaction in the ventral eye region (LM: F1,20 = 14.52, p = 0.0011): again, *H. chestertonii* males had fewer red ommatidia ($p < 0.001$), but females did not differ ($p = 0.17$). Dorsal eyeshine did not differ ($p > 0.11$; Fig. S9C).

**Hybrids have intermediate visual system morphology and visual acuity.** We generated first- (F1) and second-generation (F2) hybrids of *H. erato cyrbia* and *H. himera*, all of which were reared alongside the parental populations at Ikiam. For each hybrid, we quantified eye and brain morphology and measured visual acuity. Both hybrid types ($n_{F1} = 49$; $n_{F2} = 255$) had intermediate facet count, differing from *H. erato cyrbia* (F1: $p_{adj} = 0.06$; F2: $p_{adj} = 0.0012$) and *H. himera* ($p_{adj} < 0.0031$) but not from each other ($p_{adj} = 1.0$). For optic lobe volume, F1 hybrids ($n = 14$) differed from *H. erato cyrbia* ($p_{adj} = 0.03$) but not *H. himera* ($p_{adj} = 1.0$), whereas the F2s ($n = 198$) differed from *H. himera* ($p_{adj} < 0.001$) but not *H. erato cyrbia* ($p_{adj} = 1.0$; see Table S17 for individual neuropil comparisons). For visual acuity, neither hybrid type ($n_{F1}$ = 84; $n_{F2}$ = 219) differed from *H. erato cyrbia* (F1: posthoc marginal means estimate (β) = -0.080, 95% highest priority density (HPD): [-0.177, 0.022]; F2: β = -0.013, 95% HPD: [-0.089, 0.061]), but both differed from *H. himera* (F1: β = 0.147, 95% HPD: [0.046, 0.243]; F2: β = 0.079, 95% HPD: [0.007, 0.152]; Fig. 4B).

**Peripheral and neural structures jointly determine visual acuity (continued)**. We used a Bayesian framework to show that F2 hybrid visual acuity was positively influenced by a *facet count x optic lobe* interaction (see main text). To examine the role of individual neuropils, we initially tested a global model containing facet count and all measured visual neuropils. In this global model, the 95% CIs for all structural traits contained zero, but the lamina (LAM) and accessory medulla (aME), together with facet count, displayed the strongest directional tendencies (i.e. the positive 80% CIs were largely positive; Fig. S9A). To better investigate how these two neuropils and eye morphology together shape visual perception, we reduced our focus to these specific traits (an interactive model spanning all neuropils is computationally complex and difficult to interpret). The LAM is the first optic neuropil to receive peripheral photoreceptor input, whereas the aME is primarily associated with circadian control in insects (2); in locusts, principal neurons of the aME respond only weakly to stationary light and fail to track moving stimuli (3), making a direct, standalone role in visual acuity unlikely. Consequently, we constructed a targeted model evaluating the independent and interactive effects of facet count and LAM, while retaining aME as a covariate (in addition to tibia length and central brain volume). Mirroring the whole optic lobe analysis, the *facet count x LAM* interaction was positive in its central estimate, though the 95% credible interval slightly overlapped zero (β = 0.15, 95% CI: [-0.01, 0.31]). The conditional main effects of facet count (β = 0.11, 95% CI: [-0.82, 1.05]) and LAM (β = -0.18, 95% CI: [-0.98, 0.64]) both overlapped zero (Fig. S9B); as with the main optic lobe model, these represent baseline slopes evaluated at zero on the log10 scale—well outside our biological range—safely defaulting to our conservative prior framework [*b ~ Normal(0, 0.5)*]. The 95% CI for the aME similarly encompassed zero (β = 0.23, 95% CI: [-0.07, 0.53]). These results further support our finding that visual acuity is mediated by the interaction between eye and brain structures but specifically identify the lamina as a key neural trait. The accessory medulla may also play an indirect role (see discussion).

### Supplementary tables

**Table S1.** Sample overview of wild *H. erato* populations surveyed for eye morphology. Populations defined as in (4).

| population | year(s) collected | count | |
|---|---|---|---|
| | | m | f |
| *H. chestertonii* | 2022 | 12 | 7 |
| *H. erato venus* | 2003, 22-23 | 7 | 6 |
| *H. erato cyrbia* (North/South) | 2017, 22-23 | 41 | 25 |
| *H. erato hydara* (Panama) | 2003 | 3 | 7 |
| *H. erato hydara* (Colombia) | 1998 | 4 | 4 |
| *H. erato hydara* (French Guiana) | 2001 | 9 | 5 |
| *H. erato erato* | 2006, 07 | 15 | - |
| *H. erato notabilis* | 2009-2010 | 15 | 14 |
| *H. erato lativitta* (Ecuador) | 2009-10, 18-19, 22-23 | 41 | 23 |
| *H. erato lativitta* (Colombia) | 2017 | 3 | 3 |
| *H. himera* | 2022-23 | 25 | 17 |
| *H. erato favorinus* | 2002 | 13 | 2 |
| *H. erato emma* | 2002 | 9 | 4 |

**Table S2.** Sample overview of wild vs insectary-reared comparisons. In Ecuador, wild *H. erato cyrbia* and *H. himera* were restricted to locations used for previous brain morphology reports ((5); forests near Balsas/Buenaventura and Vilcabamba, Ecuador, respectively). Wild *H. erato lativitta* was restricted to forests near Tena, Ecuador. These three locations were used to establish the insectary populations reared under common gardens conditions at Universidad Regional Amazónica Ikiam in Tena, Ecuador. In Colombia, *H. chestertonii* (from Montañitas and El Saladito, Colombia) and *H. erato venus* (Buenaventura and La Barra) were reared under common garden conditions at José Celestino Mutis Experimental Field Station (Universidad del Rosario), near La Vega, Colombia. Prior work also reared *chestertonii-venus* in the same location (1).

| **location** | **species** | **type** | **trait** | **count** | |
|---|---|---|---|---|---|
| | | | | **m** | **f** |
| Ecuador | *H. erato cyrbia* | wild | eye | 27 | 15 |
| | *H. erato lativitta* | wild | eye | 29 | 17 |
| | *H. himera* | wild | eye | 25 | 17 |
| | *H. erato cyrbia* | insectary | eye | 14 | 19 |
| | *H. erato lativitta* | insectary | eye | 19 | 12 |
| | *H. himera* | insectary | eye | 19 | 19 |
| | *H. erato cyrbia* | wild | brain* | 8 | 8 |
| | *H. erato lativitta* | wild | brain | 7 | 7 |
| | *H. himera* | wild | brain* | 8 | 8 |
| | *H. erato cyrbia* | insectary | brain | 8 | 7 |
| | *H. erato lativitta* | insectary | brain | 6 | 6 |
| | *H. himera* | insectary | brain | 8 | 8 |
| | *H. erato cyrbia* | insectary | visual acuity | 26 | 22 |
| | *H. erato lativitta* | insectary | visual acuity | 25 | 23 |
| | *H. himera* | insectary | visual acuity | 24 | 25 |
| | *H. erato cyrbia* | insectary | eyeshine | 7 | 6 |
| | *H. erato lativitta* | insectary | eyeshine | 6 | 6 |
| | *H. himera* | insectary | eyeshine | 6 | 5 |
| | F1 *cyrbia-himera* hybrid | insectary | eye | 29 | 20 |
| | F2 *cyrbia-himera* hybrid | insectary | eye | 119 | 136 |
| | F1 *cyrbia-himera* hybrid | insectary | brain | 8 | 6 |
| | F2 *cyrbia-himera* hybrid | insectary | brain | 94 | 104 |
| | F1 *cyrbia-himera* hybrid | insectary | visual acuity | 34 | 50 |
| | F2 *cyrbia-himera* hybrid | insectary | visual acuity | 117 | 102 |
| Colombia | *H. chestertonii* | wild | eye | 12 | 7 |
| | *H. erato venus* | wild | eye | 7 | 6 |
| | *H. chestertonii* | insectary | eye | 15 | 1 |
| | *H. erato venus* | insectary | eye | 15 | 3 |
| | *H. chestertonii* | insectary | brain† | 7 | 10 |
| | *H. erato venus* | insectary | brain† | 5 | 13 |
| | *H. chestertonii* | insectary | eyeshine | 6 | 8 |
| | *H. erato venus* | insectary | eyeshine | 6 | 4 |
| | *H. chestertonii* | insectary | visual acuity | 35 | 32 |
| | *H. erato venus* | insectary | visual acuity | 33 | 31 |

** data from (5); † data from (1)*

**Table S3**. Parameter estimates from linear mixed models exploring eye morphology across wild *H. erato* populations. Sampling location included as a random effect.

| response variable | fixed effect | Wald $\chi^2$ | df | p-value |
|---|---|---|---|---|
| log10 (facet count) | population | 52.79 | 12 | **<0.001** |
| | sex | 82.16 | 1 | **<0.001** |
| | log10 (tibia length) | 204.82 | 1 | **<0.001** |
| log10 (corneal area) | population | 104.85 | 12 | **<0.001** |
| | sex | 125.45 | 1 | **<0.001** |
| | log10 (tibia length) | 308.16 | 1 | **<0.001** |

**Table S4**. Parameter estimates from linear models exploring brain morphology in wild *H. erato cyrbia*, *H. erato lativitta*, and *H. himera*. Neuropil volumes for *H. erato cyrbia* and *H. himera* from (5).

| response variable | fixed effect | F-value | Df (numerator) | Df (residual) | p-value |
|---|---|---|---|---|---|
| log10 (OL) | population | 52.29 | 2 | 40 | **<0.001** |
| | sex | 1.04 | 1 | 40 | 0.31 |
| | log10 (rCBR) | 97.19 | 1 | 40 | **<0.001** |
| log10 (LAM) | population*sex | 2.78 | 2 | 38 | 0.074 |
| | population | 49.05 | 2 | 38 | **<0.001** |
| | sex | 2.43 | 1 | 38 | 0.12 |
| | log10 (rCBR) | 13.13 | 1 | 38 | **<0.001** |
| log10 (ME) | population | 29.35 | 2 | 41 | **<0.001** |
| | sex | 0.91 | 1 | 41 | 0.35 |
| | log10 (rCBR) | 157.09 | 1 | 41 | **<0.001** |
| log10 (LOB) | population*sex | 7.42 | 2 | 39 | **0.0018** |
| | population | 65.88 | 2 | 39 | **<0.001** |
| | sex | 2.97 | 1 | 39 | 0.092 |
| | log10 (rCBR) | 53.58 | 1 | 39 | **<0.001** |
| log10 (LOP) | population*sex | 2.28 | 2 | 39 | 0.074 |
| | population | 47.93 | 2 | 39 | **<0.001** |
| | sex | 6.32 | 1 | 39 | **0.017** |
| | log10 (rCBR) | 53.13 | 1 | 39 | **<0.001** |
| log10 (vLOB) | population | 5.80 | 2 | 41 | **0.006** |
| | sex | 13.52 | 1 | 41 | **<0.001** |
| | log10 (rCBR) | 23.43 | 1 | 41 | **<0.001** |
| log10 (aME) | population | 3.63 | 2 | 41 | **0.036** |
| | sex | 3.70 | 1 | 41 | 0.061 |
| | log10 (rCBR) | 12.43 | 1 | 41 | **0.001** |
| log10 (AOTU) | population | 8.42 | 2 | 41 | **<0.001** |
| | sex | 1.02 | 1 | 41 | 0.32 |
| | log10 (rCBR) | 7.47 | 1 | 41 | **0.009** |

LAM = lamina, ME = medulla, LOB = lobula, LOP = lobula plate, vLOB = ventral lobe of the lobula, aME = accessory medulla, AOTU = anterior optic tubercule. OL = LAM + ME + LOB + LOP + vLOB + aME.

**Table S5.** Pairwise comparisons of brain morphology for wild *H. erato cyrbia*, *H. erato lativitta*, and *H. himera*, accounting for sex and central brain volume. Neuropil volumes for *H. erato cyrbia* and *H. himera* from (5). P-values are Bonferroni-adjusted ($p_{adj}$).

| trait | | comparison | estimate | SE | t ratio | $p_{adj}$ |
|---|---|---|---|---|---|---|
| OL | | *H. erato cyrbia - H. erato lativitta* | -0.0737 | 0.0125 | -5.887 | **<0.001** |
| | | *H. erato cyrbia - H. himera* | 0.0534 | 0.0117 | 4.549 | **<0.001** |
| | | *H. erato lativitta - H. himera* | 0.1271 | 0.0124 | 10.224 | **<0.001** |
| LAM | *M* | *H. erato cyrbia - H. erato lativitta* | -0.133 | 0.03 | -4.46 | 0.0002 |
| | | *H. erato cyrbia - H. himera* | 0.045 | 0.029 | 1.541 | 0.395 |
| | | *H. erato lativitta - H. himera* | 0.178 | 0.023 | 5.966 | **<0.001** |
| | *F* | *H. erato cyrbia - H. erato lativitta* | -0.236 | 0.0322 | -7.334 | **<0.001** |
| | | *H. erato cyrbia - H. himera* | -0.006 | 0.0301 | -0.2 | 1 |
| | | *H. erato lativitta - H. himera* | 0.230 | 0.0326 | 7.05 | **<0.001** |
| ME | | *H. erato cyrbia - H. erato lativitta* | -0.0368 | 0.0125 | -2.932 | **0.016** |
| | | *H. erato cyrbia - H. himera* | 0.0561 | 0.0116 | 4.838 | **<0.001** |
| | | *H. erato lativitta - H. himera* | 0.0929 | 0.0124 | 7.491 | **<0.001** |
| LOB | *M* | *H. erato cyrbia - H. erato lativitta* | -0.0732 | 0.0217 | -3.366 | **0.005** |
| | | *H. erato cyrbia - H. himera* | 0.0525 | 0.021 | 2.499 | 0.0503 |
| | | *H. erato lativitta - H. himera* | 0.1257 | 0.0217 | 5.795 | **<0.001** |
| | *F* | *H. erato cyrbia - H. erato lativitta* | -0.155 | 0.0233 | -6.661 | **<0.001** |
| | | *H. erato cyrbia - H. himera* | 0.0927 | 0.021 | 4.421 | **0.0002** |
| | | *H. erato lativitta - H. himera* | 0.2476 | 0.0234 | 10.597 | **<0.001** |
| *LOP* | *M* | *H. erato cyrbia - H. erato lativitta* | -0.083 | 0.0229 | -3.608 | 0.003 |
| | | *H. erato cyrbia - H. himera* | 0.083 | 0.0222 | 3.747 | 0.002 |
| | | *H. erato lativitta - H. himera* | 0.166 | 0.0229 | 7.247 | **<0.001** |
| | *F* | *H. erato cyrbia - H. erato lativitta* | -0.144 | 0.0245 | -5.862 | **<0.001** |
| | | *H. erato cyrbia - H. himera* | 0.016 | 0.0221 | 0.71 | 1 |
| | | *H. erato lativitta - H. himera* | 0.159 | 0.0246 | 6.472 | **<0.001** |
| vLOB | | *H. erato cyrbia - H. erato lativitta* | -0.0707 | 0.0271 | -2.611 | **0.038** |
| | | *H. erato cyrbia - H. himera* | 0.0171 | 0.025 | 0.685 | 1 |
| | | *H. erato lativitta - H. himera* | 0.0878 | 0.0268 | 3.282 | **0.006** |
| aME | | *H. erato cyrbia - H. erato lativitta* | -0.0952 | 0.0433 | -2.2 | 0.100 |
| | | *H. erato cyrbia - H. himera* | -0.0964 | 0.04 | -2.41 | 0.062 |
| | | *H. erato lativitta - H. himera* | -0.0012 | 0.0428 | -0.028 | 1 |
| AOTU | | *H. erato cyrbia - H. erato lativitta* | -0.1234 | 0.0366 | -3.367 | **0.005** |
| | | *H. erato cyrbia - H. himera* | 0.0158 | 0.0339 | 0.466 | 1 |
| | | *H. erato lativitta - H. himera* | 0.1392 | 0.0362 | 3.843 | **0.001** |

LAM = lamina, ME = medulla, LOB = lobula, LOP = lobula plate, vLOB = ventral lobe of the lobula, aME = accessory medulla, AOTU = anterior optic tubercle. OL = LAM + ME + LOB + LOP + vLOB + aME.

**Table S6**. Parameter estimates from linear models exploring eye morphology in insectary-reared *H. erato cyrbia, H. erato lativitta,* and *H. himera.*

| response variable | fixed effect | F-value | Df (numerator) | Df (residual) | p-value |
|---|---|---|---|---|---|
| log10 (facet count) | population | 37.54 | 2 | 97 | **<0.001** |
| | sex | 36.76 | 1 | 97 | **<0.001** |
| | log10 (tibia length) | 3.88 | 1 | 97 | 0.052 |
| log10 (corneal area) | population | 89.72 | 2 | 97 | **<0.001** |
| | sex | 63.59 | 1 | 97 | **<0.001** |
| | log10 (tibia length) | 26.91 | 1 | 97 | **<0.001** |

**Table S7**. Parameter estimates from linear models exploring brain morphology in insectary-reared *H. erato cyrbia*, *H. erato lativitta*, and *H. himera*.

| response variable | fixed effect | F-value | Df (numerator) | Df (residual) | p-value |
|---|---|---|---|---|---|
| log10 (OL) | population * sex | 5.22 | 2 | 32 | **0.011** |
| | population | 16.65 | 2 | 32 | **<0.001** |
| | sex | 4.16 | 1 | 32 | **0.049** |
| | log10 (rCBR) | 280.05 | 1 | 32 | **<0.001** |
| log10 (LAM) | population * sex | 3.55 | 2 | 32 | **0.040** |
| | population | 8.85 | 2 | 32 | **<0.001** |
| | sex | 0.52 | 1 | 32 | 0.48 |
| | log10 (rCBR) | 73.19 | 1 | 32 | **<0.001** |
| log10 (ME) | population * sex | 8.44 | 2 | 36 | **<0.001** |
| | population | 15.89 | 2 | 36 | **<0.001** |
| | sex | 6.72 | 1 | 36 | **0.014** |
| | log10 (rCBR) | 381.84 | 1 | 36 | **<0.001** |
| log10 (LOB) | population * sex | 5.20 | 2 | 36 | **0.010** |
| | population | 8.80 | 2 | 36 | **<0.001** |
| | sex | 1.73 | 1 | 36 | 0.19 |
| | log10 (rCBR) | 363.56 | 1 | 36 | **<0.001** |
| log10 (LOP | population | 2.23 | 2 | 38 | 0.12 |
| | sex | 4.07 | 1 | 38 | 0.051 |
| | log10 (rCBR) | 289.59 | 1 | 38 | **<0.001** |
| log10 (vLOB) | population | 1.58 | 2 | 38 | 0.22 |
| | sex | 7.40 | 1 | 38 | **0.009** |
| | log10 (rCBR) | 177.95 | 1 | 38 | **<0.001** |
| log10 (aME) | population | 0.47 | 2 | 38 | 0.62 |
| | sex | 0.71 | 1 | 38 | 0.40 |
| | log10 (rCBR) | 48.25 | 1 | 38 | **<0.001** |
| log10 (AOTU) | population | 0.33 | 2 | 38 | 0.72 |
| | sex | 0.07 | 1 | 38 | 0.79 |
| | log10 (rCBR) | 110.41 | 1 | 38 | **<0.001** |

LAM = lamina, ME = medulla, LOB = lobula, LOP = lobula plate, vLOB = ventral lobe of the lobula, aME = accessory medulla, AOTU = anterior optic tubercule. OL = LAM + ME + LOB + LOP + vLOB + aME.

**Table S8.** Pairwise comparisons of brain morphology for insectary-reared *H. erato cyrbia*, *H. erato lativitta*, and *H. himera*, accounting for sex and central brain volume. P-values are Bonferroni-adjusted.

| trait | | comparison | estimate | SE | t ratio | $p_{adj}$ |
|---|---|---|---|---|---|---|
| OL | M | *H. erato cyrbia - H. erato lativitta* | 0.0174 | 0.0198 | 0.878 | 1 |
| | | *H. erato cyrbia - H. himera* | 0.0989 | 0.019 | 5.212 | **<0.001** |
| | | *H. erato lativitta - H. himera* | 0.0815 | 0.0189 | 4.326 | **0.0004** |
| | F | *H. erato cyrbia - H. erato lativitta* | -0.0651 | 0.0219 | -2.977 | **0.0165** |
| | | *H. erato cyrbia - H. himera* | 0.0078 | 0.0193 | 0.404 | 1 |
| | | *H. erato lativitta - H. himera* | 0.0729 | 0.0193 | 3.77 | **0.002** |
| LAM | M | *H. erato cyrbia - H. erato lativitta* | 0.004 | 0.037 | 0.108 | 1 |
| | | *H. erato cyrbia - H. himera* | 0.1001 | 0.0354 | 2.825 | **0.0243** |
| | | *H. erato lativitta - H. himera* | 0.0961 | 0.0352 | 2.731 | **0.0305** |
| | F | *H. erato cyrbia - H. erato lativitta* | -0.1481 | 0.0408 | -3.627 | **0.003** |
| | | *H. erato cyrbia - H. himera* | -0.0159 | 0.036 | -0.441 | 1 |
| | | *H. erato lativitta - H. himera* | 0.1322 | 0.0361 | 3.662 | **0.0027** |
| ME | M | *H. erato cyrbia - H. erato lativitta* | 0.0319 | 0.0169 | 1.89 | 0.2004 |
| | | *H. erato cyrbia - H. himera* | 0.1051 | 0.017 | 6.169 | **<0.001** |
| | | *H. erato lativitta - H. himera* | 0.0732 | 0.0166 | 4.415 | **0.0003** |
| | F | *H. erato cyrbia - H. erato lativitta* | -0.0482 | 0.0179 | -2.694 | **0.032** |
| | | *H. erato cyrbia - H. himera* | 0.0074 | 0.0159 | 0.465 | 1 |
| | | *H. erato lativitta - H. himera* | 0.0556 | 0.0165 | 3.371 | **0.0054** |
| LOB | M | *H. erato cyrbia - H. erato lativitta* | 0.041 | 0.02 | 2.05 | 0.1432 |
| | | *H. erato cyrbia - H. himera* | 0.0966 | 0.0202 | 4.776 | **<0.001** |
| | | *H. erato lativitta - H. himera* | 0.0556 | 0.0197 | 2.823 | **0.0231** |
| | F | *H. erato cyrbia - H. erato lativitta* | -0.0418 | 0.0212 | -1.968 | 0.1703 |
| | | *H. erato cyrbia - H. himera* | 0.0094 | 0.0189 | 0.498 | 1 |
| | | *H. erato lativitta - H. himera* | 0.0512 | 0.0196 | 2.616 | **0.0388** |
| LOP | | *H. erato cyrbia - H. erato lativitta* | -0.0032 | 0.0182 | -0.174 | 1 |
| | | *H. erato cyrbia - H. himera* | 0.03083 | 0.0172 | 1.79 | 0.2445 |
| | | *H. erato lativitta - H. himera* | 0.034 | 0.0184 | 1.843 | 0.2194 |
| vLOB | | *H. erato cyrbia - H. erato lativitta* | 0.0108 | 0.0245 | 0.441 | 1 |
| | | *H. erato cyrbia - H. himera* | 0.0402 | 0.0232 | 1.731 | 0.2746 |
| | | *H. erato lativitta - H. himera* | 0.0294 | 0.0248 | 1.182 | 0.7342 |
| aME | | *H. erato cyrbia - H. erato lativitta* | 0.0401 | 0.0421 | 0.952 | 1 |
| | | *H. erato cyrbia - H. himera* | 0.0247 | 0.0399 | 0.62 | 1 |
| | | *H. erato lativitta - H. himera* | -0.0154 | 0.0427 | -0.36 | 1 |
| AOTU | | *H. erato cyrbia - H. erato lativitta* | 0.01531 | 0.0247 | 0.619 | 1 |
| | | *H. erato cyrbia - H. himera* | -0.0045 | 0.0234 | -0.192 | 1 |
| | | *H. erato lativitta - H. himera* | -0.0198 | 0.0251 | -0.79 | 1 |

LAM = lamina, ME = medulla, LOB = lobula, LOP = lobula plate, vLOB = ventral lobe of the lobula, aME = accessory medulla, AOTU = anterior optic tubercule. OL = LAM + ME + LOB + LOP + vLOB + aME.

**Table S9.** Samples used for windowed $F_{ST}$ genome-wide analyses. ENA = European Nucleotide Archive. NCBI = National Center for Biotechnology Information.

| sample id | species | source |
|---|---|---|
| HE.cyrbia.14N014 | *Heliconius erato cyrbia* | publicly available at ENA |
| HE.cyrbia.15N016 | *Heliconius erato cyrbia* | publicly available at ENA |
| HE.cyrbia.15N017 | *Heliconius erato cyrbia* | publicly available at ENA |
| HE.cyrbia.CAM040752 | *Heliconius erato cyrbia* | publicly available at ENA |
| HE.cyrbia_ERR3789821 | *Heliconius erato cyrbia* | publicly available at ENA |
| HE.cyrbia_ERR3789822 | *Heliconius erato cyrbia* | publicly available at ENA |
| HE.cyrbia_ERR3789823 | *Heliconius erato cyrbia* | publicly available at ENA |
| HE.cyrbia_ERR3789824 | *Heliconius erato cyrbia* | publicly available at ENA |
| HE.cyrbia_ERR3789825 | *Heliconius erato cyrbia* | publicly available at ENA |
| HE.cyrbia_ERR3789826 | *Heliconius erato cyrbia* | publicly available at ENA |
| HE.cyrbia_ERR3789827 | *Heliconius erato cyrbia* | publicly available at ENA |
| HE.cyrbia_LMU_20023 | *Heliconius erato cyrbia* | this study |
| HE.cyrbia_LMU_20292 | *Heliconius erato cyrbia* | this study |
| HE.cyrbia_LMU_20295 | *Heliconius erato cyrbia* | this study |
| HE.cyrbia_LMU_20337 | *Heliconius erato cyrbia* | this study |
| HE.himera001 | *Heliconius himera* | publicly available at NCBI |
| HE.himera002 | *Heliconius himera* | publicly available at NCBI |
| HE.himera003 | *Heliconius himera* | publicly available at NCBI |
| HE.himera030 | *Heliconius himera* | publicly available at NCBI |
| HE.himera_LMU_20024 | *Heliconius himera* | this study |
| HE.himera_LMU_20025 | *Heliconius himera* | this study |
| HE.himera_LMU_20026 | *Heliconius himera* | this study |
| HE.himera_LMU_20027 | *Heliconius himera* | this study |
| HE.himera_LMU_20028 | *Heliconius himera* | this study |
| HE.himera_LMU_20030 | *Heliconius himera* | this study |
| HE.himera_LMU_20155 | *Heliconius himera* | this study |
| HE.himera_LMU_20194 | *Heliconius himera* | this study |
| HE.himera_LMU_20233 | *Heliconius himera* | this study |
| HE.himera_LMU_20328 | *Heliconius himera* | this study |
| HE.himera_LMU_20372 | *Heliconius himera* | this study |
| HE.himera_LMU_20890 | *Heliconius himera* | this study |
| HE.himera_LMU_20901 | *Heliconius himera* | this study |
| HE.himera_LMU_20917 | *Heliconius himera* | this study |
| HE.himera_LMU_20997 | *Heliconius himera* | this study |
| HE.himera_LMU_20998 | *Heliconius himera* | this study |
| HE.himera_LMU_21004 | *Heliconius himera* | this study |
| HE.himera_LMU_21005 | *Heliconius himera* | this study |
| HE.himera_LMU_21006 | *Heliconius himera* | this study |
| HE.himera_SRR4032038 | *Heliconius himera* | publicly available at ENA |
| HE.himera_SRR4032039 | *Heliconius himera* | publicly available at ENA |
| HE.himera_SRR4032040 | *Heliconius himera* | publicly available at ENA |
| HE.himera_SRR4032041 | *Heliconius himera* | publicly available at ENA |
| HE.himera_SRR6313531 | *Heliconius himera* | publicly available at ENA |
| HE.himera_SRR6313532 | *Heliconius himera* | publicly available at ENA |

| | | |
|---|---|---|
| HE.himera_SRR6313541 | *Heliconius himera* | publicly available at ENA |
| HE.himera_SRR6313542 | *Heliconius himera* | publicly available at ENA |
| HE.lativitta_ERR3781215 | *Heliconius erato lativitta* | publicly available at ENA |
| HE.lativitta_ERR3789812 | *Heliconius erato lativitta* | publicly available at ENA |
| HE.lativitta_ERR3789813 | *Heliconius erato lativitta* | publicly available at ENA |
| HE.lativitta_ERR3789814 | *Heliconius erato lativitta* | publicly available at ENA |
| HE.lativitta_ERR3789815 | *Heliconius erato lativitta* | publicly available at ENA |
| HE.lativitta_ERR3789816 | *Heliconius erato lativitta* | publicly available at ENA |
| HE.lativitta_ERR3789817 | *Heliconius erato lativitta* | publicly available at ENA |
| HE.lativitta_ERR3789818 | *Heliconius erato lativitta* | publicly available at ENA |
| HE.lativitta_ERR3789819 | *Heliconius erato lativitta* | publicly available at ENA |
| HE.lativitta_ERR3789820 | *Heliconius erato lativitta* | publicly available at ENA |
| HE.lativitta_ERR3789830 | *Heliconius erato lativitta* | publicly available at ENA |
| HE.lativitta_ERR3789831 | *Heliconius erato lativitta* | publicly available at ENA |
| HE.lativitta_ERR3789846 | *Heliconius erato lativitta* | publicly available at ENA |
| HE.lativitta_ERR3789847 | *Heliconius erato lativitta* | publicly available at ENA |
| HE.lativitta_ERR3789848 | *Heliconius erato lativitta* | publicly available at ENA |
| HE.lativitta_ERR3789849 | *Heliconius erato lativitta* | publicly available at ENA |
| HE.lativitta_ERR3790131 | *Heliconius erato lativitta* | publicly available at ENA |
| HE.lativitta_ERR3790132 | *Heliconius erato lativitta* | publicly available at ENA |
| HE.lativitta_ERR3790133 | *Heliconius erato lativitta* | publicly available at ENA |
| HE.lativitta_ERR3790134 | *Heliconius erato lativitta* | publicly available at ENA |
| HE.lativitta_ERR3790135 | *Heliconius erato lativitta* | publicly available at ENA |
| HE.lativitta_ERR9907861 | *Heliconius erato lativitta* | publicly available at ENA |
| HE.lativitta_ERR9907867 | *Heliconius erato lativitta* | publicly available at ENA |
| HE.lativitta_LMU_20376 | *Heliconius erato lativitta* | this study |
| HE.lativitta_SRR4032044 | *Heliconius erato lativitta* | publicly available at NCBI |
| HE.lativitta_SRR4032045 | *Heliconius erato lativitta* | publicly available at NCBI |
| HE.lativitta_SRR4032046 | *Heliconius erato lativitta* | publicly available at NCBI |

**Table S10**. $P_{ST}$—$F_{ST}$ comparisons for eye morphology between *H. erato cyrbia* and *H. himera*. P-values calculated as the proportion of the $F_{ST}$ distribution that is above each $P_{ST}$ value; values above the 95th percentile ($P_{95}$) of the $F_{ST}$ distribution are interpreted as an indication of selection. False discovery rate (FDR) correction applied for multiple testing.

| trait | $F_{ST}$ ($P_{95}$) | c/$h^2$ | $P_{ST}$ | lower 95% CI | upper 95% CI | FDR p-value |
|---|---|---|---|---|---|---|
| facet count | 0.8306 | 0.33 | 0.800 | 0.515 | 0.904 | 0.081 |
| | | 0.67 | 0.891 | 0.709 | 0.950 | **0.017** |
| | | 1.00 | 0.924 | 0.789 | 0.967 | **0.006** |
| | | 1.33 | 0.942 | 0.826 | 0.974 | **0.003** |
| | | 1.50 | 0.948 | 0.837 | 0.978 | **0.003** |
| | | 2.00 | 0.960 | 0.881 | 0.984 | **0.002** |
| | | 3.00 | 0.973 | 0.908 | 0.989 | **0.001** |
| | | 4.00 | 0.980 | 0.940 | 0.991 | **0.001** |
| corneal area | 0.8306 | 0.33 | 0.917 | 0.863 | 0.950 | **0.007** |
| | | 0.67 | 0.958 | 0.927 | 0.974 | **0.002** |
| | | 1.00 | 0.971 | 0.949 | 0.983 | **0.001** |
| | | 1.33 | 0.978 | 0.961 | 0.987 | **0.001** |
| | | 1.50 | 0.981 | 0.967 | 0.989 | **0.001** |
| | | 2.00 | 0.985 | 0.974 | 0.991 | **0.001** |
| | | 3.00 | 0.990 | 0.983 | 0.994 | **0.001** |
| | | 4.00 | 0.993 | 0.987 | 0.996 | **0.001** |

**Table S11**. $P_{ST}$—$F_{ST}$ comparisons for eye morphology between *H. erato lativitta* and *H. himera*. P-values calculated as the proportion of the $F_{ST}$ distribution that is above each $P_{ST}$ value; values above the 95th percentile ($P_{95}$) of the $F_{ST}$ distribution are interpreted as an indication of selection. False discovery rate (FDR) correction applied for multiple testing.

| trait | $F_{ST}$ ($P_{95}$) | c/$h^2$ | $P_{ST}$ | lower 95% CI | upper 95% CI | FDR p-value |
|---|---|---|---|---|---|---|
| facet count | 0.5254 | 0.33 | 0.791 | 0.608 | 0.879 | **0.000** |
| | | 0.67 | 0.885 | 0.760 | 0.937 | **0.000** |
| | | 1.00 | 0.920 | 0.827 | 0.958 | **0.000** |
| | | 1.33 | 0.938 | 0.858 | 0.967 | **0.000** |
| | | 1.50 | 0.945 | 0.875 | 0.972 | **0.000** |
| | | 2.00 | 0.958 | 0.906 | 0.978 | **0.000** |
| | | 3.00 | 0.972 | 0.938 | 0.986 | **0.000** |
| | | 4.00 | 0.979 | 0.953 | 0.989 | **0.000** |
| corneal area | 0.5254 | 0.33 | 0.845 | 0.739 | 0.913 | **0.000** |
| | | 0.67 | 0.917 | 0.844 | 0.954 | **0.000** |
| | | 1.00 | 0.943 | 0.890 | 0.968 | **0.000** |
| | | 1.33 | 0.957 | 0.923 | 0.976 | **0.000** |
| | | 1.50 | 0.961 | 0.924 | 0.979 | **0.000** |
| | | 2.00 | 0.971 | 0.939 | 0.984 | **0.000** |
| | | 3.00 | 0.980 | 0.962 | 0.989 | **0.000** |
| | | 4.00 | 0.985 | 0.969 | 0.992 | **0.000** |

**Table S12**. $P_{ST}$—$F_{ST}$ comparisons for eye morphology between *H. erato cyrbia* and *H. erato lativitta*. P-values calculated as the proportion of the $F_{ST}$ distribution that is above each $P_{ST}$ value; values above the 95<sup>th</sup> percentile ($P_{95}$) of the $F_{ST}$ distribution are interpreted as an indication of selection. False discovery rate (FDR) correction applied for multiple testing.

| trait | $F_{ST}$ ($P_{95}$) | c/$h^2$ | $P_{ST}$ | lower 95% CI | upper 95% CI | FDR p-value |
|---|---|---|---|---|---|---|
| facet count | 0.5460 | 0.33 | 0.132 | 0.000 | 0.557 | 0.998 |
| | | 0.67 | 0.235 | 0.001 | 0.726 | 0.998 |
| | | 1.00 | 0.315 | 0.002 | 0.793 | 0.741 |
| | | 1.33 | 0.379 | 0.001 | 0.833 | 0.591 |
| | | 1.50 | 0.408 | 0.003 | 0.842 | 0.591 |
| | | 2.00 | 0.479 | 0.002 | 0.885 | 0.475 |
| | | 3.00 | 0.579 | 0.002 | 0.924 | 0.288 |
| | | 4.00 | 0.647 | 0.010 | 0.936 | 0.288 |
| corneal area | 0.5460 | 0.33 | 0.017 | 0.000 | 0.489 | 0.998 |
| | | 0.67 | 0.034 | 0.001 | 0.648 | 0.998 |
| | | 1.00 | 0.050 | 0.001 | 0.742 | 0.998 |
| | | 1.33 | 0.066 | 0.001 | 0.779 | 0.998 |
| | | 1.50 | 0.074 | 0.002 | 0.801 | 0.998 |
| | | 2.00 | 0.096 | 0.001 | 0.864 | 0.998 |
| | | 3.00 | 0.137 | 0.002 | 0.909 | 0.998 |
| | | 4.00 | 0.175 | 0.004 | 0.915 | 0.998 |

**Table S13**. $P_{ST}$—$F_{ST}$ comparisons for brain morphology between *H. erato cyrbia* and *H. himera*. P-values calculated as the proportion of the $F_{ST}$ distribution that is above each $P_{ST}$ value; values above the 95[th] percentile ($P_{95}$) of the $F_{ST}$ distribution are interpreted as an indication of selection. False discovery rate (FDR) correction applied for multiple testing.

| trait | $F_{ST}$ ($P_{95}$) | c/$h^2$ | $P_{ST}$ | lower 95% CI | upper 95% CI | FDR p-value |
|---|---|---|---|---|---|---|
| OL | 0.8306 | 0.33 | 0.713 | 0.361 | 0.889 | 0.400 |
| | | 0.67 | 0.834 | 0.545 | 0.940 | 0.163 |
| | | 1.00 | 0.883 | 0.624 | 0.962 | 0.081 |
| | | 1.33 | 0.909 | 0.708 | 0.968 | **0.048** |
| | | 1.50 | 0.919 | 0.730 | 0.974 | **0.039** |
| | | 2.00 | 0.938 | 0.794 | 0.981 | **0.020** |
| | | 3.00 | 0.958 | 0.825 | 0.987 | **0.015** |
| | | 4.00 | 0.968 | 0.887 | 0.990 | **0.015** |
| LAM | 0.8306 | 0.33 | 0.295 | 0.002 | 0.758 | 0.999 |
| | | 0.67 | 0.459 | 0.007 | 0.864 | 0.972 |
| | | 1.00 | 0.559 | 0.005 | 0.895 | 0.739 |
| | | 1.33 | 0.628 | 0.004 | 0.917 | 0.596 |
| | | 1.50 | 0.655 | 0.006 | 0.931 | 0.526 |
| | | 2.00 | 0.717 | 0.012 | 0.952 | 0.400 |
| | | 3.00 | 0.792 | 0.012 | 0.965 | 0.253 |
| | | 4.00 | 0.835 | 0.021 | 0.973 | 0.163 |
| ME | 0.8306 | 0.33 | 0.804 | 0.598 | 0.914 | 0.240 |
| | | 0.67 | 0.893 | 0.766 | 0.959 | 0.073 |
| | | 1.00 | 0.926 | 0.828 | 0.969 | **0.034** |
| | | 1.33 | 0.943 | 0.861 | 0.977 | **0.020** |
| | | 1.50 | 0.949 | 0.882 | 0.979 | **0.020** |
| | | 2.00 | 0.961 | 0.905 | 0.985 | **0.015** |
| | | 3.00 | 0.974 | 0.931 | 0.990 | **0.015** |
| | | 4.00 | 0.980 | 0.948 | 0.992 | **0.015** |
| LOB | 0.8306 | 0.33 | 0.710 | 0.366 | 0.878 | 0.400 |
| | | 0.67 | 0.833 | 0.531 | 0.940 | 0.163 |
| | | 1.00 | 0.881 | 0.649 | 0.958 | 0.081 |
| | | 1.33 | 0.908 | 0.704 | 0.967 | **0.049** |
| | | 1.50 | 0.918 | 0.722 | 0.972 | **0.039** |
| | | 2.00 | 0.937 | 0.799 | 0.981 | **0.020** |
| | | 3.00 | 0.957 | 0.864 | 0.987 | **0.015** |
| | | 4.00 | 0.967 | 0.870 | 0.989 | **0.015** |
| LOP | 0.8306 | 0.33 | 0.383 | 0.011 | 0.737 | 0.999 |
| | | 0.67 | 0.558 | 0.018 | 0.852 | 0.739 |
| | | 1.00 | 0.653 | 0.049 | 0.888 | 0.526 |
| | | 1.33 | 0.714 | 0.054 | 0.915 | 0.400 |
| | | 1.50 | 0.738 | 0.041 | 0.927 | 0.373 |
| | | 2.00 | 0.790 | 0.043 | 0.947 | 0.253 |
| | | 3.00 | 0.849 | 0.093 | 0.961 | 0.133 |
| | | 4.00 | 0.883 | 0.099 | 0.972 | 0.081 |
| vLOB | 0.8306 | 0.33 | 0.251 | 0.001 | 0.675 | 0.999 |
| | | 0.67 | 0.405 | 0.003 | 0.797 | 0.999 |
| | | 1.00 | 0.504 | 0.005 | 0.850 | 0.873 |
| | | 1.33 | 0.575 | 0.005 | 0.879 | 0.728 |
| | | 1.50 | 0.604 | 0.010 | 0.901 | 0.648 |
| | | 2.00 | 0.670 | 0.004 | 0.920 | 0.500 |
| | | 3.00 | 0.753 | 0.006 | 0.945 | 0.343 |
| | | 4.00 | 0.803 | 0.025 | 0.964 | 0.240 |
| aME | 0.8306 | 0.33 | 0.094 | 0.000 | 0.625 | 0.999 |
| | | 0.67 | 0.174 | 0.000 | 0.808 | 0.999 |
| | | 1.00 | 0.239 | 0.001 | 0.843 | 0.999 |

| | | | | | | |
|---|---|---|---|---|---|---|
| | | 1.33 | 0.295 | 0.001 | 0.884 | 0.999 |
| | | 1.50 | 0.320 | 0.002 | 0.896 | 0.999 |
| | | 2.00 | 0.386 | 0.002 | 0.919 | 0.999 |
| | | 3.00 | 0.485 | 0.003 | 0.941 | 0.917 |
| | | 4.00 | 0.557 | 0.005 | 0.953 | 0.739 |
| AOTU | 0.8306 | 0.33 | 0.010 | 0.000 | 0.528 | 0.999 |
| | | 0.67 | 0.020 | 0.001 | 0.695 | 0.999 |
| | | 1.00 | 0.030 | 0.001 | 0.756 | 0.999 |
| | | 1.33 | 0.040 | 0.001 | 0.813 | 0.999 |
| | | 1.50 | 0.045 | 0.001 | 0.850 | 0.999 |
| | | 2.00 | 0.059 | 0.002 | 0.855 | 0.999 |
| | | 3.00 | 0.085 | 0.001 | 0.914 | 0.999 |
| | | 4.00 | 0.111 | 0.003 | 0.931 | 0.999 |

LAM = lamina, ME = medulla, LOB = lobula, LOP = lobula plate, vLOB = ventral lobe of the lobula, aME = accessory medulla, AOTU = anterior optic tubercule. OL = LAM + ME + LOB + LOP + vLOB + aME.

**Table S14**. $P_{ST}$—$F_{ST}$ comparisons for brain morphology between *H. erato lativitta* and *H. himera*. P-values calculated as the proportion of the $F_{ST}$ distribution that is above each $P_{ST}$ value; values above the 95th percentile ($P_{95}$) of the $F_{ST}$ distribution are interpreted as an indication of selection. False discovery rate (FDR) correction applied for multiple testing.

| trait | $F_{ST}$ ($P_{95}$) | c/$h^2$ | $P_{ST}$ | lower 95% CI | upper 95% CI | FDR p-value |
|---|---|---|---|---|---|---|
| OL | 0.5254 | 0.33 | 0.809 | 0.605 | 0.923 | **0.004** |
| | | 0.67 | 0.896 | 0.758 | 0.961 | **0.000** |
| | | 1.00 | 0.928 | 0.823 | 0.973 | **0.000** |
| | | 1.33 | 0.945 | 0.850 | 0.979 | **0.000** |
| | | 1.50 | 0.950 | 0.872 | 0.983 | **0.000** |
| | | 2.00 | 0.962 | 0.895 | 0.986 | **0.000** |
| | | 3.00 | 0.975 | 0.930 | 0.991 | **0.000** |
| | | 4.00 | 0.981 | 0.945 | 0.993 | **0.000** |
| LAM | 0.5254 | 0.33 | 0.798 | 0.531 | 0.939 | 0.782 |
| | | 0.67 | 0.889 | 0.687 | 0.963 | 0.209 |
| | | 1.00 | 0.923 | 0.775 | 0.976 | **0.045** |
| | | 1.33 | 0.941 | 0.818 | 0.982 | **0.015** |
| | | 1.50 | 0.947 | 0.844 | 0.984 | **0.009** |
| | | 2.00 | 0.960 | 0.876 | 0.989 | **0.004** |
| | | 3.00 | 0.973 | 0.919 | 0.991 | **0.000** |
| | | 4.00 | 0.980 | 0.935 | 0.994 | **0.000** |
| ME | 0.5254 | 0.33 | 0.817 | 0.684 | 0.910 | **0.000** |
| | | 0.67 | 0.900 | 0.829 | 0.952 | **0.000** |
| | | 1.00 | 0.931 | 0.877 | 0.967 | **0.000** |
| | | 1.33 | 0.947 | 0.905 | 0.975 | **0.000** |
| | | 1.50 | 0.953 | 0.917 | 0.978 | **0.000** |
| | | 2.00 | 0.964 | 0.933 | 0.985 | **0.000** |
| | | 3.00 | 0.976 | 0.956 | 0.989 | **0.000** |
| | | 4.00 | 0.982 | 0.966 | 0.992 | **0.000** |
| LOB | 0.5254 | 0.33 | 0.740 | 0.450 | 0.886 | **0.004** |
| | | 0.67 | 0.852 | 0.618 | 0.942 | **0.000** |
| | | 1.00 | 0.896 | 0.707 | 0.961 | **0.000** |
| | | 1.33 | 0.920 | 0.758 | 0.970 | **0.000** |
| | | 1.50 | 0.928 | 0.769 | 0.972 | **0.000** |
| | | 2.00 | 0.945 | 0.842 | 0.981 | **0.000** |
| | | 3.00 | 0.963 | 0.890 | 0.987 | **0.000** |
| | | 4.00 | 0.972 | 0.908 | 0.990 | **0.000** |
| LOP | 0.5254 | 0.33 | 0.374 | 0.006 | 0.732 | 0.442 |
| | | 0.67 | 0.548 | 0.017 | 0.842 | **0.045** |
| | | 1.00 | 0.644 | 0.033 | 0.894 | **0.009** |
| | | 1.33 | 0.707 | 0.029 | 0.915 | **0.004** |
| | | 1.50 | 0.731 | 0.032 | 0.926 | **0.002** |
| | | 2.00 | 0.784 | 0.049 | 0.938 | **0.000** |
| | | 3.00 | 0.845 | 0.106 | 0.962 | **0.000** |
| | | 4.00 | 0.879 | 0.080 | 0.973 | **0.000** |
| vLOB | 0.5254 | 0.33 | 0.283 | 0.003 | 0.744 | 0.931 |
| | | 0.67 | 0.445 | 0.007 | 0.865 | 0.379 |
| | | 1.00 | 0.545 | 0.005 | 0.889 | 0.113 |
| | | 1.33 | 0.615 | 0.007 | 0.912 | **0.037** |
| | | 1.50 | 0.643 | 0.011 | 0.922 | **0.024** |
| | | 2.00 | 0.706 | 0.013 | 0.945 | **0.007** |

| | | | | | | |
|---|---|---|---|---|---|---|
| | | 3.00 | 0.782 | 0.019 | 0.963 | **0.002** |
| | | 4.00 | 0.827 | 0.027 | 0.971 | **0.000** |
| aME | 0.5254 | 0.33 | 0.000 | 0.000 | 0.532 | 0.999 |
| | | 0.67 | 0.000 | 0.000 | 0.718 | 0.999 |
| | | 1.00 | 0.001 | 0.001 | 0.785 | 0.963 |
| | | 1.33 | 0.001 | 0.001 | 0.837 | 0.782 |
| | | 1.50 | 0.001 | 0.002 | 0.831 | 0.691 |
| | | 2.00 | 0.001 | 0.001 | 0.874 | 0.435 |
| | | 3.00 | 0.002 | 0.003 | 0.923 | 0.147 |
| | | 4.00 | 0.003 | 0.002 | 0.940 | **0.045** |
| AOTU | 0.5254 | 0.33 | 0.156 | 0.001 | 0.687 | 0.999 |
| | | 0.67 | 0.274 | 0.001 | 0.820 | 0.999 |
| | | 1.00 | 0.360 | 0.003 | 0.856 | 0.999 |
| | | 1.33 | 0.428 | 0.002 | 0.888 | 0.999 |
| | | 1.50 | 0.457 | 0.004 | 0.905 | 0.999 |
| | | 2.00 | 0.529 | 0.002 | 0.924 | 0.999 |
| | | 3.00 | 0.628 | 0.006 | 0.949 | 0.999 |
| | | 4.00 | 0.692 | 0.011 | 0.960 | 0.999 |

LAM = lamina, ME = medulla, LOB = lobula, LOP = lobula plate, vLOB = ventral lobe of the lobula, aME = accessory medulla, AOTU = anterior optic tubercule. OL = LAM + ME + LOB + LOP + vLOB + aME.

**Table S15**. $P_{ST}$—$F_{ST}$ comparisons for brain morphology between *H. erato cyrbia* and *H. erato lativitta*. P-values calculated as the proportion of the $F_{ST}$ distribution that is above each $P_{ST}$ value; values above the 95th percentile ($P_{95}$) of the $F_{ST}$ distribution are interpreted as an indication of selection. False discovery rate (FDR) correction applied for multiple testing.

| trait | $F_{ST}$ ($P_{95}$) | $c/h^2$ | $P_{ST}$ | lower 95% CI | upper 95% CI | FDR p-value |
|---|---|---|---|---|---|---|
| OL | 0.5460 | 0.33 | 0.141 | 0.000 | 0.622 | 0.999 |
| | | 0.67 | 0.250 | 0.001 | 0.764 | 0.985 |
| | | 1.00 | 0.333 | 0.002 | 0.816 | 0.663 |
| | | 1.33 | 0.399 | 0.004 | 0.852 | 0.525 |
| | | 1.50 | 0.428 | 0.004 | 0.873 | 0.485 |
| | | 2.00 | 0.499 | 0.003 | 0.895 | 0.358 |
| | | 3.00 | 0.599 | 0.006 | 0.939 | 0.190 |
| | | 4.00 | 0.666 | 0.010 | 0.945 | 0.152 |
| LAM | 0.5460 | 0.33 | 0.437 | 0.039 | 0.760 | 0.485 |
| | | 0.67 | 0.612 | 0.068 | 0.870 | 0.190 |
| | | 1.00 | 0.702 | 0.063 | 0.904 | 0.112 |
| | | 1.33 | 0.758 | 0.142 | 0.929 | 0.064 |
| | | 1.50 | 0.779 | 0.146 | 0.938 | 0.064 |
| | | 2.00 | 0.825 | 0.209 | 0.949 | 0.054 |
| | | 3.00 | 0.876 | 0.295 | 0.966 | **0.022** |
| | | 4.00 | 0.904 | 0.345 | 0.976 | **0.015** |
| ME | 0.5460 | 0.33 | 0.012 | 0.000 | 0.457 | 0.999 |
| | | 0.67 | 0.025 | 0.000 | 0.663 | 0.999 |
| | | 1.00 | 0.037 | 0.001 | 0.744 | 0.999 |
| | | 1.33 | 0.048 | 0.001 | 0.789 | 0.999 |
| | | 1.50 | 0.054 | 0.001 | 0.803 | 0.999 |
| | | 2.00 | 0.071 | 0.001 | 0.870 | 0.999 |
| | | 3.00 | 0.102 | 0.002 | 0.896 | 0.999 |
| | | 4.00 | 0.132 | 0.001 | 0.911 | 0.999 |
| LOB | 0.5460 | 0.33 | 0.019 | 0.000 | 0.493 | 0.999 |
| | | 0.67 | 0.037 | 0.001 | 0.642 | 0.999 |
| | | 1.00 | 0.054 | 0.000 | 0.741 | 0.999 |
| | | 1.33 | 0.071 | 0.001 | 0.771 | 0.999 |
| | | 1.50 | 0.080 | 0.001 | 0.805 | 0.999 |
| | | 2.00 | 0.103 | 0.001 | 0.867 | 0.999 |
| | | 3.00 | 0.147 | 0.002 | 0.900 | 0.999 |
| | | 4.00 | 0.187 | 0.003 | 0.930 | 0.999 |
| LOP | 0.5460 | 0.33 | 0.000 | 0.000 | 0.511 | 0.999 |
| | | 0.67 | 0.000 | 0.000 | 0.625 | 0.999 |
| | | 1.00 | 0.000 | 0.000 | 0.710 | 0.999 |
| | | 1.33 | 0.000 | 0.001 | 0.788 | 0.999 |
| | | 1.50 | 0.000 | 0.001 | 0.803 | 0.999 |
| | | 2.00 | 0.000 | 0.002 | 0.845 | 0.999 |
| | | 3.00 | 0.000 | 0.001 | 0.902 | 0.999 |
| | | 4.00 | 0.000 | 0.001 | 0.921 | 0.999 |
| vLOB | 0.5460 | 0.33 | 0.015 | 0.000 | 0.531 | 0.999 |
| | | 0.67 | 0.030 | 0.000 | 0.707 | 0.999 |
| | | 1.00 | 0.044 | 0.001 | 0.794 | 0.999 |
| | | 1.33 | 0.058 | 0.001 | 0.828 | 0.999 |
| | | 1.50 | 0.065 | 0.002 | 0.846 | 0.999 |
| | | 2.00 | 0.085 | 0.002 | 0.879 | 0.999 |

| | | | | | | |
|---|---|---|---|---|---|---|
| | | 3.00 | 0.122 | 0.001 | 0.921 | 0.999 |
| | | 4.00 | 0.157 | 0.002 | 0.949 | 0.999 |
| aME | 0.5460 | 0.33 | 0.055 | 0.000 | 0.557 | 0.999 |
| | | 0.67 | 0.105 | 0.000 | 0.736 | 0.999 |
| | | 1.00 | 0.149 | 0.001 | 0.811 | 0.999 |
| | | 1.33 | 0.189 | 0.001 | 0.839 | 0.999 |
| | | 1.50 | 0.208 | 0.002 | 0.875 | 0.999 |
| | | 2.00 | 0.259 | 0.003 | 0.895 | 0.961 |
| | | 3.00 | 0.344 | 0.002 | 0.923 | 0.660 |
| | | 4.00 | 0.412 | 0.003 | 0.935 | 0.523 |
| AOTU | 0.5460 | 0.33 | 0.110 | 0.000 | 0.567 | 0.999 |
| | | 0.67 | 0.200 | 0.001 | 0.740 | 0.999 |
| | | 1.00 | 0.272 | 0.002 | 0.830 | 0.920 |
| | | 1.33 | 0.332 | 0.001 | 0.852 | 0.663 |
| | | 1.50 | 0.359 | 0.002 | 0.874 | 0.639 |
| | | 2.00 | 0.427 | 0.003 | 0.895 | 0.485 |
| | | 3.00 | 0.528 | 0.004 | 0.927 | 0.313 |
| | | 4.00 | 0.599 | 0.005 | 0.945 | 0.190 |

LAM = lamina, ME = medulla, LOB = lobula, LOP = lobula plate, vLOB = ventral lobe of the lobula, aME = accessory medulla, AOTU = anterior optic tubercule. OL = LAM + ME + LOB + LOP + vLOB + aME.

**Table S16**. Parameter estimates from linear mixed models exploring eye morphology in insectary-reared *H. erato cyrbia, H. himera*, and their hybrids. Family identity included as a random effect.

| response variable | fixed effect | Wald χ2 | df | p-value |
|---|---|---|---|---|
| log10 (facet count) | population | 40.36 | 3 | **<0.001** |
| | sex | 151.30 | 1 | **<0.001** |
| | log10 (tibia length) | 69.52 | 1 | **<0.001** |
| log10 (corneal area) | population | 148.36 | 3 | **<0.001** |
| | sex | 333.69 | 1 | **<0.001** |
| | log10 (tibia length) | 183.15 | 1 | **<0.001** |

**Table S17**. Parameter estimates from linear mixed models exploring brain morphology in insectary-reared *H. erato cyrbia, H. himera*, and their hybrids. Family identity included as a random effect.

| response variable | fixed effect | Wald χ2 | df | p-value |
|---|---|---|---|---|
| log10 (OL) | population * sex | 8.08 | 3 | **0.044** |
| | population | 31.22 | 3 | **<0.001** |
| | sex | 2.90 | 1 | 0.088 |
| | log10 (rCBR) | 1342.17 | 1 | **<0.001** |
| log10 (LAM) | population | 13.04 | 3 | **0.005** |
| | sex | 8.01 | 1 | **0.005** |
| | log10 (rCBR) | 531.03 | 1 | **<0.001** |
| log10 (ME) | population * sex | 11.41 | 3 | **0.009** |
| | population | 18.38 | 3 | **<0.001** |
| | sex | 6.63 | 1 | **0.010** |
| | log10 (rCBR) | 1366.88 | 1 | **<0.001** |
| log10 (LOB) | population * sex | 12.82 | 3 | **0.005** |
| | population | 18.19 | 3 | **<0.001** |
| | sex | 2.69 | 1 | 0.101 |
| | log10 (rCBR) | 1400.13 | 1 | **<0.001** |
| log10 (LOP) | population | 10.74 | 3 | **0.013** |
| | sex | 5.64 | 1 | **0.018** |
| | log10 (rCBR) | 575.26 | 1 | **<0.001** |
| log10 (vLOB) | population | 3.65 | 3 | 0.30 |
| | sex | 57.81 | 1 | **<0.001** |
| | log10 (rCBR) | 355.71 | 1 | **<0.001** |
| log10 (aME) | population | 9.96 | 3 | **0.019** |
| | sex | 1.02 | 1 | 0.31 |
| | log10 (rCBR) | 87.56 | 1 | **<0.001** |
| log10 (AOTU) | population | 16.79 | 3 | **<0.001** |
| | sex | 11.57 | 1 | **<0.001** |
| | log10 (rCBR) | 460.36 | 1 | **<0.001** |

LAM = lamina, ME = medulla, LOB = lobula, LOP = lobula plate, vLOB = ventral lobe of the lobula, aME = accessory medulla, AOTU = anterior optic tubercule. OL = LAM + ME + LOB + LOP + vLOB + aME.

**Table S18.** Pairwise comparisons of brain morphology for insectary-reared *H. erato cyrbia*, *H. himera*, and their hybrids, accounting for sex, central brain volume, and family identity (as a random effect). P-values are Bonferroni-adjusted.

| trait | | comparison | estimate | SE | t ratio | $p_{adj}$ |
|---|---|---|---|---|---|---|
| OL | M | *H. erato cyrbia* - F1 hybrids | 0.06059 | 0.0166 | 3.66 | **0.0022** |
| | | *H. erato cyrbia* - F2 hybrids | 0.03382 | 0.013 | 2.611 | 0.0606 |
| | | *H. erato cyrbia - H. himera* | 0.07924 | 0.0168 | 4.725 | **<0.001** |
| | | F1 hybrids - F2 hybrids | -0.02677 | 0.0128 | -2.095 | 0.2339 |
| | | F1 hybrids - *H. himera* | 0.01865 | 0.0165 | 1.128 | 1 |
| | | F2 hybrids - *H. himera* | 0.04542 | 0.0122 | 3.718 | **0.0018** |
| | F | *H. erato cyrbia* - F1 hybrids | 0.00978 | 0.019 | 0.515 | 1 |
| | | *H. erato cyrbia* - F2 hybrids | -0.01794 | 0.0139 | -1.289 | 1 |
| | | *H. erato cyrbia - H. himera* | 0.02102 | 0.0181 | 1.16 | 1 |
| | | F1 hybrids - F2 hybrids | -0.02772 | 0.0146 | -1.897 | 0.3659 |
| | | F1 hybrids - *H. himera* | 0.01124 | 0.0186 | 0.604 | 1 |
| | | F2 hybrids - *H. himera* | 0.03896 | 0.0135 | 2.884 | **0.0274** |
| LAM | | *H. erato cyrbia* - F1 hybrids | 0.03374 | 0.022 | 1.537 | 0.7644 |
| | | *H. erato cyrbia* - F2 hybrids | -0.00672 | 0.0165 | -0.406 | 1 |
| | | *H. erato cyrbia - H. himera* | 0.04079 | 0.0206 | 1.978 | 0.2976 |
| | | F1 hybrids - F2 hybrids | -0.04046 | 0.0173 | -2.334 | 0.1432 |
| | | F1 hybrids - *H. himera* | 0.00705 | 0.0215 | 0.328 | 1 |
| | | F2 hybrids - *H. himera* | 0.0475 | 0.0157 | 3.027 | **0.0182** |
| ME | M | *H. erato cyrbia* - F1 hybrids | -0.010803 | 0.0144 | -0.749 | 1 |
| | | *H. erato cyrbia* - F2 hybrids | -0.063714 | 0.0186 | -3.424 | **0.0049** |
| | | *H. erato cyrbia - H. himera* | 0.029013 | 0.0186 | 1.56 | 0.7264 |
| | | F1 hybrids - F2 hybrids | -0.052911 | 0.014 | -3.789 | **0.0013** |
| | | F1 hybrids - *H. himera* | 0.039816 | 0.0136 | 2.924 | **0.0242** |
| | | F2 hybrids - *H. himera* | 0.092727 | 0.0184 | 5.037 | **<0.001** |
| | F | *H. erato cyrbia* - F1 hybrids | -0.009198 | 0.0165 | -0.558 | 1 |
| | | *H. erato cyrbia* - F2 hybrids | -0.000399 | 0.0207 | -0.019 | 1 |
| | | *H. erato cyrbia - H. himera* | 0.012259 | 0.0203 | 0.605 | 1 |
| | | F1 hybrids - F2 hybrids | 0.0088 | 0.0143 | 0.614 | 1 |
| | | F1 hybrids - *H. himera* | 0.021457 | 0.0136 | 1.575 | 0.7051 |
| | | F2 hybrids - *H. himera* | 0.012658 | 0.0185 | 0.683 | 1 |
| LOB | M | *H. erato cyrbia* - F1 hybrids | -0.01015 | 0.0153 | -0.663 | 1 |
| | | *H. erato cyrbia* - F2 hybrids | -0.07299 | 0.0197 | -3.712 | **0.0019** |
| | | *H. erato cyrbia - H. himera* | 0.02551 | 0.0196 | 1.298 | 1 |
| | | F1 hybrids - F2 hybrids | -0.06283 | 0.0147 | -4.262 | **0.0002** |
| | | F1 hybrids - *H. himera* | 0.03566 | 0.0144 | 2.48 | 0.0865 |
| | | F2 hybrids - *H. himera* | 0.09849 | 0.0194 | 5.082 | **<0.001** |
| | F | *H. erato cyrbia* - F1 hybrids | -0.02752 | 0.0175 | -1.573 | 0.713 |
| | | *H. erato cyrbia* - F2 hybrids | -0.02397 | 0.0219 | -1.095 | 1 |
| | | *H. erato cyrbia - H. himera* | -0.01537 | 0.0214 | -0.718 | 1 |
| | | F1 hybrids - F2 hybrids | 0.00355 | 0.0151 | 0.235 | 1 |
| | | F1 hybrids - *H. himera* | 0.01215 | 0.0144 | 0.844 | 1 |
| | | F2 hybrids - *H. himera* | 0.00859 | 0.0195 | 0.44 | 1 |
| LOP | | *H. erato cyrbia* - F1 hybrids | -0.030322 | 0.0169 | -1.799 | 0.4597 |
| | | *H. erato cyrbia* - F2 hybrids | -0.06854 | 0.0214 | -3.206 | **0.0102** |

| | | | | | |
|---|---|---|---|---|---|
| | *H. erato cyrbia* - *H. himera* | -0.030115 | 0.021 | -1.434 | 0.9237 |
| | F1 hybrids - F2 hybrids | -0.038218 | 0.0153 | -2.502 | 0.08 |
| | F1 hybrids - *H. himera* | 0.000207 | 0.0147 | 0.014 | 1 |
| | F2 hybrids - *H. himera* | 0.038425 | 0.0197 | 1.949 | 0.3158 |
| vLOB | *H. erato cyrbia* - F1 hybrids | 0.02201 | 0.0291 | 0.755 | 1 |
| | *H. erato cyrbia* - F2 hybrids | -0.00383 | 0.0334 | -0.115 | 1 |
| | *H. erato cyrbia* - *H. himera* | 0.04429 | 0.033 | 1.342 | 1 |
| | F1 hybrids - F2 hybrids | -0.02585 | 0.0227 | -1.137 | 1 |
| | F1 hybrids - *H. himera* | 0.02227 | 0.022 | 1.014 | 1 |
| | F2 hybrids - *H. himera* | 0.04812 | 0.0275 | 1.751 | 0.4952 |
| aME | *H. erato cyrbia* - F1 hybrids | 0.0877 | 0.0361 | 2.427 | 0.1063 |
| | *H. erato cyrbia* - F2 hybrids | 0.0166 | 0.0464 | 0.359 | 1 |
| | *H. erato cyrbia* - *H. himera* | 0.0516 | 0.0455 | 1.133 | 1 |
| | F1 hybrids - F2 hybrids | -0.071 | 0.0333 | -2.133 | 0.2059 |
| | F1 hybrids - *H. himera* | -0.0361 | 0.0319 | -1.13 | 1 |
| | F2 hybrids - *H. himera* | 0.0349 | 0.0433 | 0.807 | 1 |
| AOTU | *H. erato cyrbia* - F1 hybrids | -0.00935 | 0.0218 | -0.429 | 1 |
| | *H. erato cyrbia* - F2 hybrids | -0.05753 | 0.0266 | -2.163 | 0.1982 |
| | *H. erato cyrbia* - *H. himera* | -0.06955 | 0.0262 | -2.657 | 0.0556 |
| | F1 hybrids - F2 hybrids | -0.04818 | 0.0187 | -2.575 | 0.067 |
| | F1 hybrids - *H. himera* | -0.0602 | 0.018 | -3.347 | **0.0065** |
| | F2 hybrids - *H. himera* | -0.01202 | 0.0237 | -0.508 | 1 |

LAM = lamina, ME = medulla, LOB = lobula, LOP = lobula plate, vLOB = ventral lobe of the lobula, aME = accessory medulla, AOTU = anterior optic tubercule. OL = LAM + ME + LOB + LOP + vLOB + aME.

**Table S19**. Parameter estimates from linear models exploring eye morphology in wild *H. erato venus* and *H. chestertonii.*

| response variable | fixed effect | F-value | Df (numerator) | Df (residual) | p-value |
|---|---|---|---|---|---|
| log10 (facet count) | population * sex | 2.93 | 1 | 27 | 0.098 |
| | population | 1.38 | 1 | 27 | 0.250 |
| | sex | 5.56 | 1 | 27 | **0.026** |
| | log10 (tibia length) | 31.85 | 1 | 27 | **<0.001** |
| log10 (corneal area) | population * sex | 3.73 | 1 | 27 | 0.064 |
| | population | 19.88 | 1 | 27 | **<0.001** |
| | sex | 3.22 | 1 | 27 | 0.083 |
| | log10 (tibia length) | 27.73 | 1 | 27 | **<0.001** |

**Table S20**. Parameter estimates from linear models exploring eye morphology in insectary-reared *H. erato venus* and *H. chestertonii.* Note: there were few female samples (*chestertonii* = 1, *venus* = 3).

| response variable | fixed effect | F-value | Df (numerator) | Df (residual) | p-value |
|---|---|---|---|---|---|
| log10 (facet count) | population | 0.271 | 1 | 30 | 0.61 |
| | sex | 29.69 | 1 | 30 | **<0.001** |
| | log10 (tibia length) | 7.25 | 1 | 30 | **0.011** |
| log10 (corneal area) | population | 56.78 | 1 | 30 | **<0.001** |
| | sex | 35.12 | 1 | 30 | **<0.001** |
| | log10 (tibia length) | 12.03 | 1 | 30 | **0.001** |

**Table S21**. Parameter estimates from linear models evaluating the relationship between facet count and corneal area for wild and insectary-reared *H. erato venus* vs the other sampled *H. erato* populations (group designated as: *H. erato venus* vs others).

| response variable | fixed effect | F-value | Df (numerator) | Df (residual) | p-value |
|---|---|---|---|---|---|
| **wild** | | | | | |
| log10 (corneal area) | group * log10 (facet count) | 0.704 | 1 | 308 | 0.474 |
| | group | 60.56 | 1 | 308 | **<0.001** |
| | log10 (facet count) | 845.57 | 1 | 308 | **<0.001** |
| | log10 (tibia length) | 34.93 | 1 | 308 | **<0.001** |
| **insectary-reared** | | | | | |
| log10 (corneal area) | group * log10 (facet count) | 2.41 | 1 | 130 | 0.122 |
| | group | 129.19 | 1 | 130 | **<0.001** |
| | log10 (facet count) | 208.88 | 1 | 130 | **<0.001** |
| | log10 (tibia length) | 14.79 | 1 | 130 | **<0.001** |

### Supplementary figures

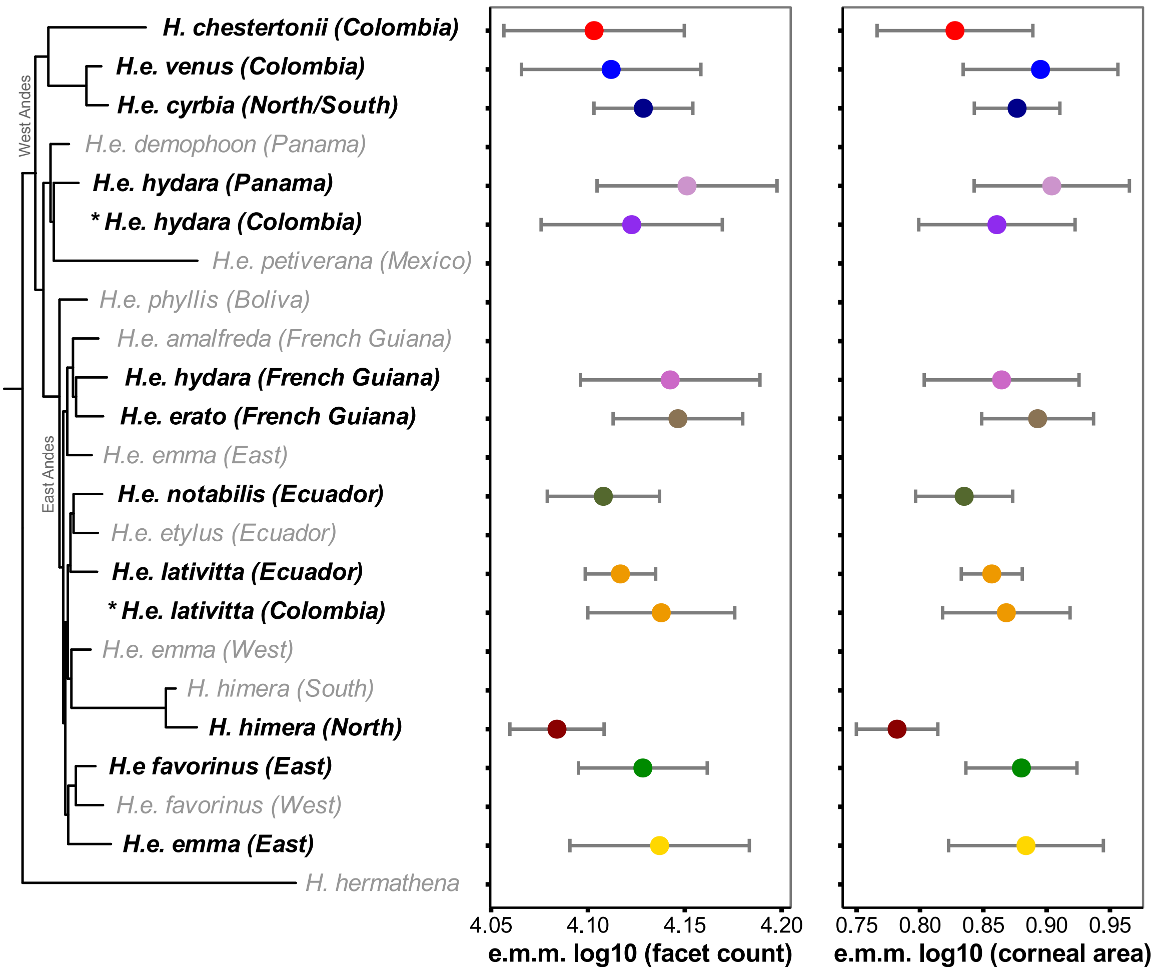


**Figure S1**. Facet count and corneal area differed across *H. erato* populations (p <0.001). Estimated marginal means (e.m.m.) account for sex, tibia length, and sampling location (as a random effect).

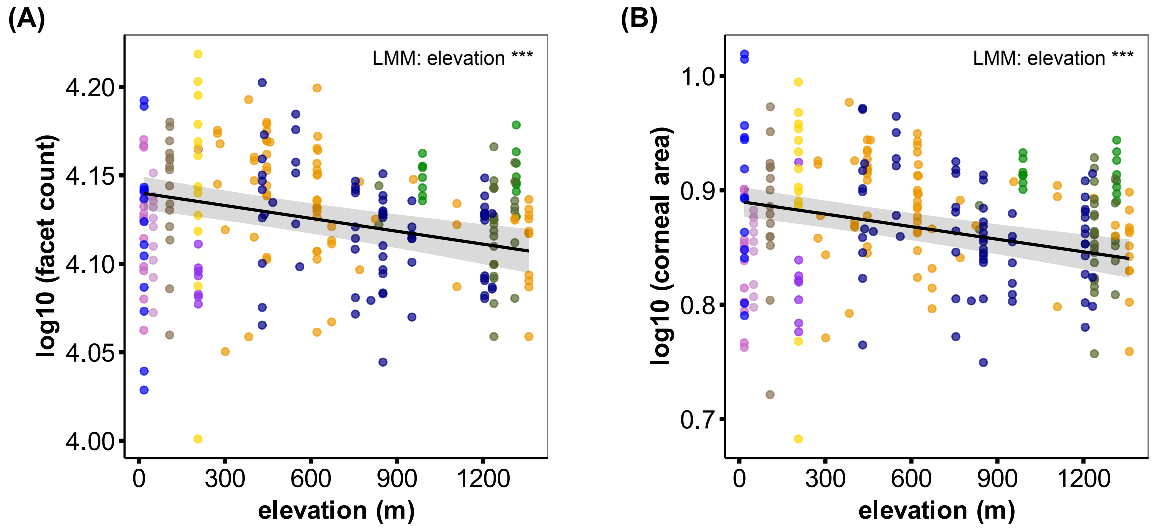


**Figure S2**. Even when excluding the two high-elevation specialists, *H. himera* and *H. chestertonii*, eye morphology was significantly influenced by elevation. Black lines represent the estimated marginal mean and 95% confidence interval (shaded ribbon) from a linear mixed model (LMM) accounting for sex, tibia length, and population (as a random effect). ***indicates p <0.001

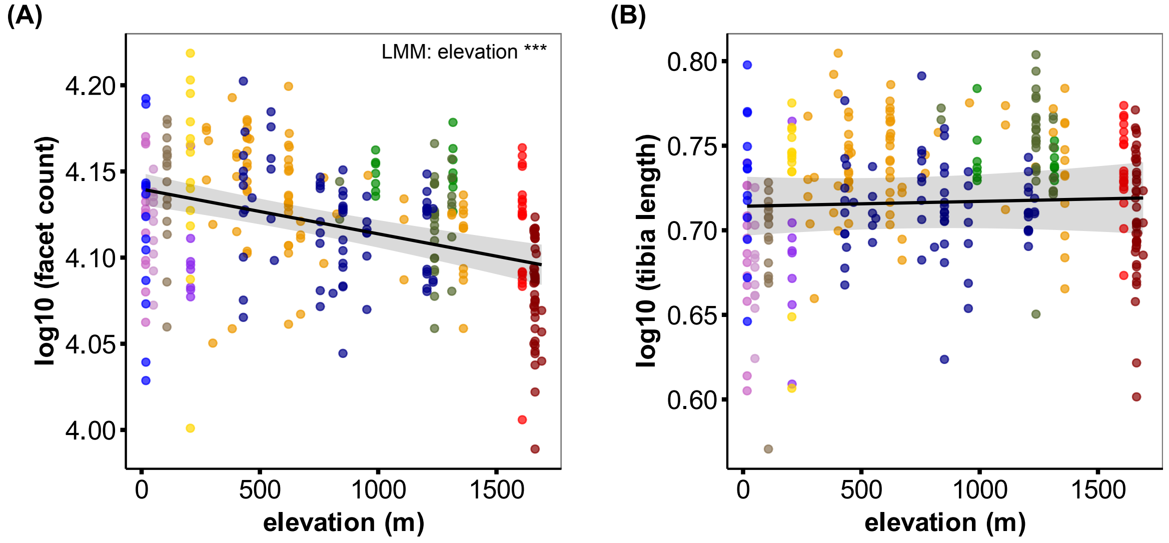


**Figure S3**. **(A)** Facet count was significantly influenced by elevation, but **(B)** tibia length (a body size proxy) did not covary with elevation. Black lines represent the estimated marginal mean and 95% confidence interval (shaded ribbon) from a linear mixed model (LMM) accounting for (A) sex, tibia length, and population (as a random effect) or (B) sex and population (as a random effect). ***indicates p <0.001.

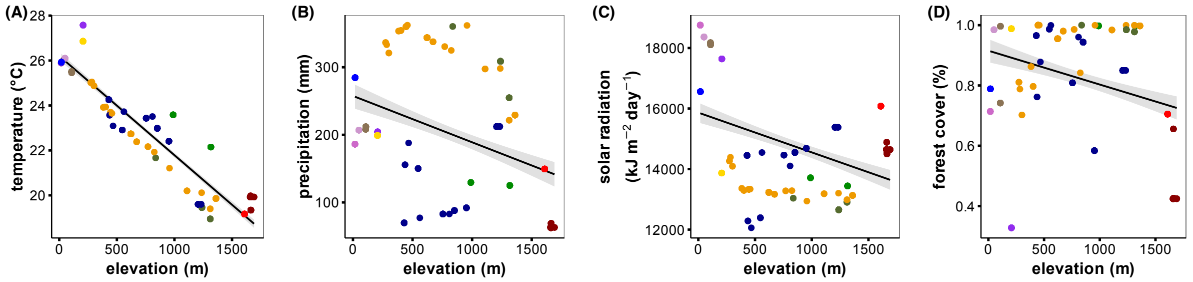


**Figure S4**. Higher elevation was associated with lower **(A)** mean annual temperature, **(B)** precipitation, and **(C)** solar radiation, and less **(D)** forest cover. Solid line represents the linear relationship and standard error (shaded ribbon).

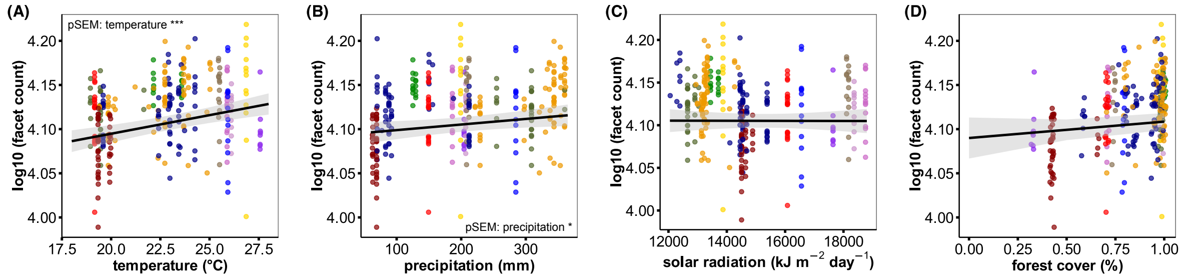


**Figure S5**. Facet count was positively influenced by **(A)** mean annual temperature and **(B)** mean annual precipitation, but **(C)** mean annual solar radiation and **(D)** forest cover had no effect. Black line represents the estimated marginal mean and 95% confidence interval (shaded ribbon) from a piecewise structural equation model (pSEM) accounting for sex, tibia length, and population (as a random effect), as well as the relationships between climatic variables (see methods for full model description). ***indicates p <0.001; *indicates p <0.05

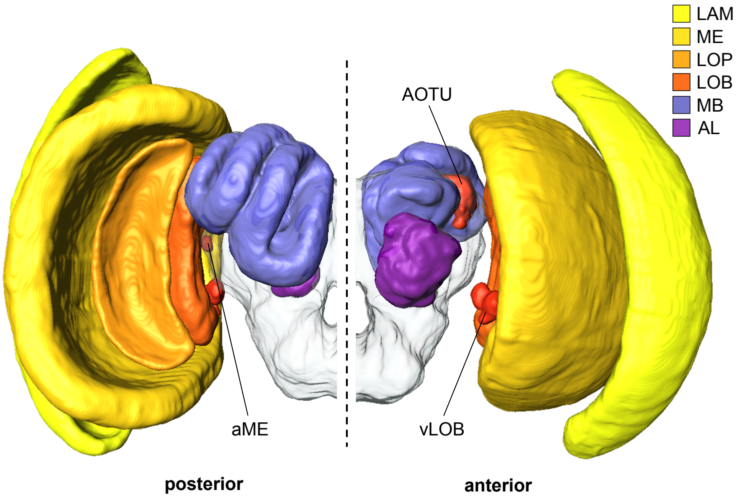


**Figure S6.** We assessed brain morphology as volumetric measurements of the visual (warm colors) and central brain neuropils. LAM = lamina, ME = medulla, LOB = lobula, LOP = lobula plate, vLOB = ventral lobe of the lobula, aME = accessory medulla, AOTU = anterior optic tubercule. MB = mushroom bodies, AL = antennal lobe. Optic lobe = LAM + ME + LOB + LOP + vLOB + aME.

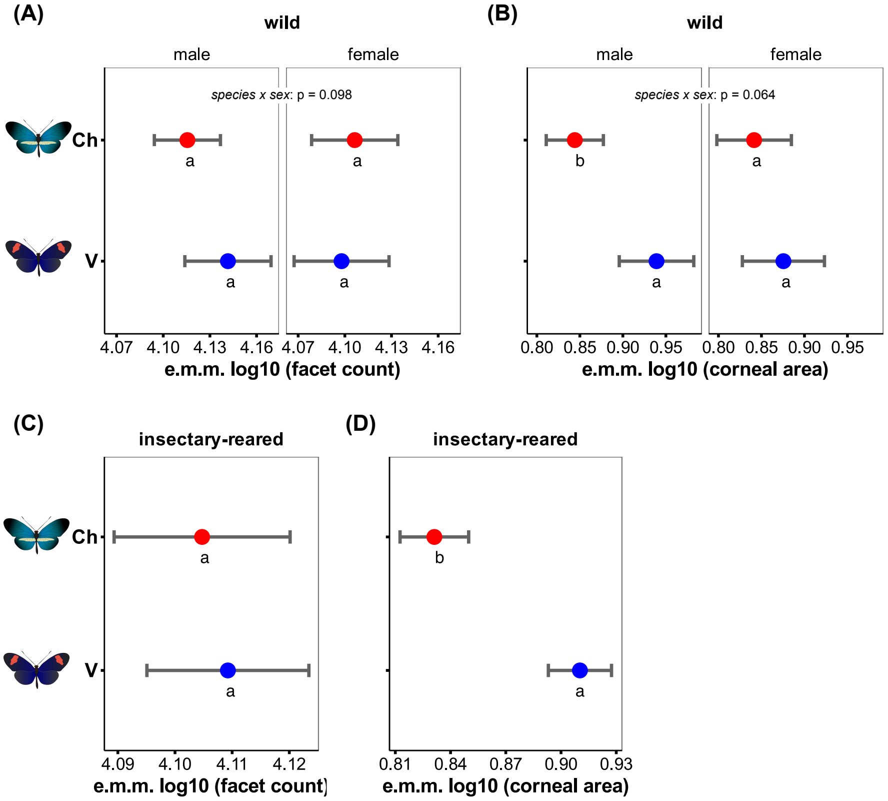


**Figure S7**. In Colombia, facet count did not differ for **(A)** wild or **(C)** insectary-reared *H. erato venus* [V] vs *H. chestertonii* [Ch], but **(B, D)** *H. erato venus* had larger corneas. Estimated marginal means (e.m.m.) account for sex and tibia length, error bars represent 95% confidence intervals, and different letters indicate significant differences (Bonferroni-adjusted p <0.05).

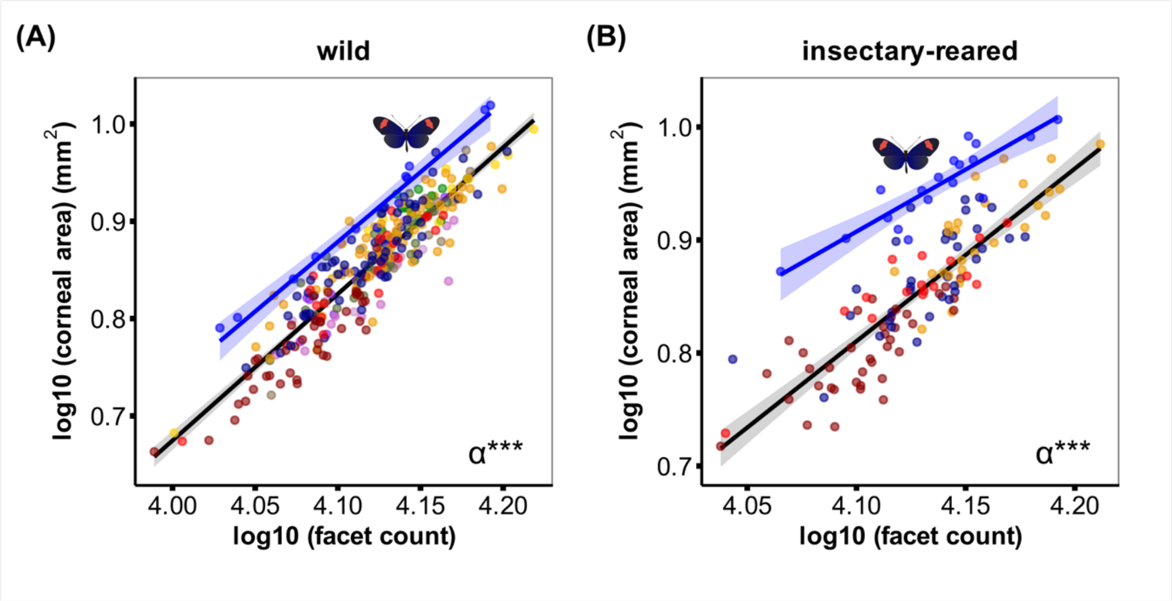


**Figure S8.** For both **(A)** wild and **(B)** insectary-reared butterflies, *H. erato venus* (blue line) displayed a significant grade shift (α) in the relationship between facet count and corneal area (slopes did not differ, p >0.13). This suggests that *H. erato venus* has evolved larger facets than the rest of the *H. erato*s sampled here. The black line represents the mean relationship for all other *eratos* (excluding *H. erato venus*), and the shaded ribbons indicate standard error. Insectary-reared butterflies from Ecuador (*H. erato cyrbia*, *H. erato lativitta*, and *H. himera*) also included in B. Population colors are the same as in figure 1. *** indicates p <0.001.

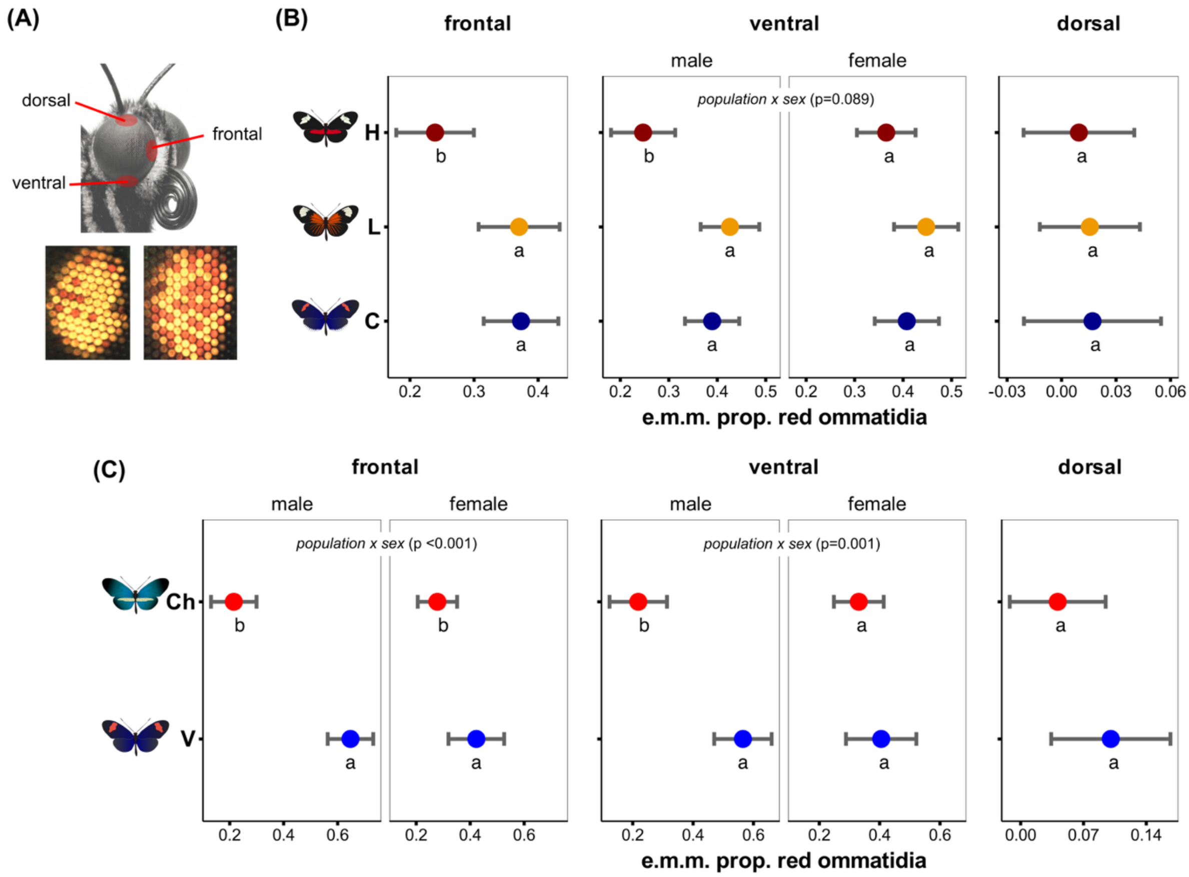


**Figure S9.** Eyeshine, a proxy for lateral filtering pigments, across **(A)** different eye regions for insectary-reared butterflies from **(B)** Ecuador (*H. himera* [H], *H. erato lativitta* [L], *H. erato cyrbia* [C]) and **(C)** Colombia (*H. chestertonii* [Ch] and *H. erato venus* [V]). Estimated marginal means (e.m.m.) account for sex, error bars represent 95% confidence intervals, and different letters indicate significant differences (Bonferroni-adjusted p <0.05). Males and females presented separately only when there was a *population x sex* interaction.

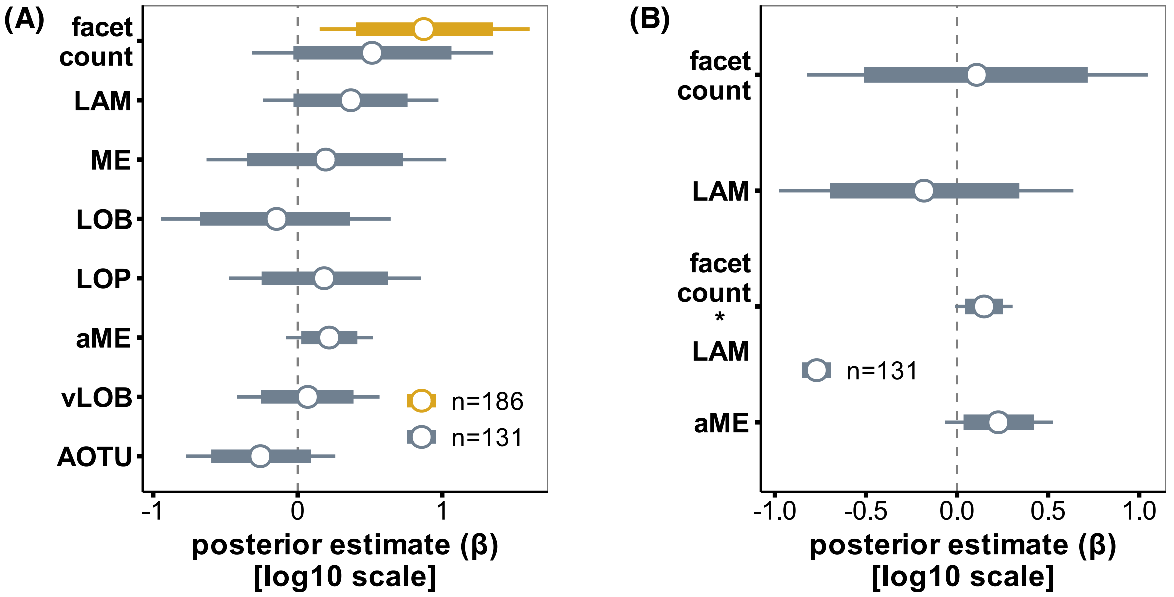


**Figure S10. (A)** For 186 F2 hybrids, facet count positively influenced visual acuity (yellow bar), whereas a smaller sub-sample of F2s with complete brain data (n=131, grey bars) suggested positive effects of facet count together with the lamina (LAM) and accessory medulla (aME). **(B)** An interactive model with only these three traits revealed a slight positive *facet count x lamina* interaction (aME was included as a covariate). Horizontal bars represent the 80% (thick) and 95% (thin) posterior intervals, with the central point indicating the posterior median from Bayesian censored models (see supplementary results and methods for complete model descriptions). LAM = lamina, ME = medulla, LOB = lobula, LOP = lobula plate, vLOB = ventral lobe of the lobula, aME = accessory medulla, AOTU = anterior optic tubercule.

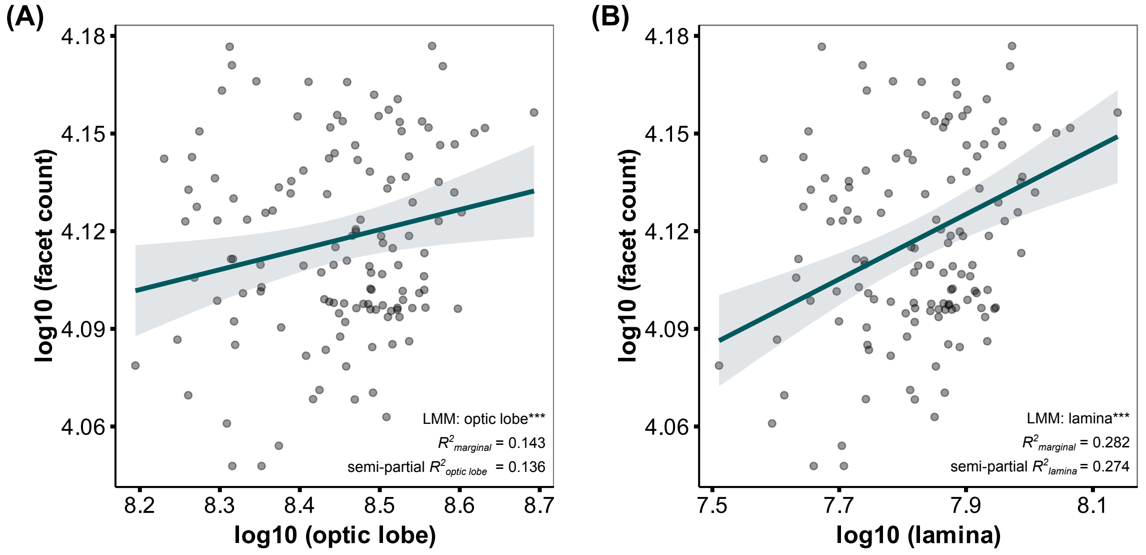


**Figure S11.** Facet count was positively correlated with **(A)** optic lobe and **(B)** lamina volume. The solid lines represent values from linear mixed models (LMM), incorporating covariation of central brain volume and controlling for family identity as a random effect. The shaded region represents the 95% confidence interval, and the points are raw observations. Marginal and semi-partial $R^2$ values obtained from *lmer()* models with *partR2()* (6). ***indicates p <0.001.
